# Targeting CREB remodels the immune microenvironment to enhance immunotherapy responses in pancreatic cancer

**DOI:** 10.64898/2025.12.04.691935

**Authors:** Siddharth Mehra, Supriya Srinivasan, Sudhakar Jinka, Varunkumar Krishnamoorthy, Vineet Kumar Gupta, Vanessa Garrido, Anna Bianchi, Andrew M Adams, Manan Patel, Haleh Amirian, Luis A Nivelo, Karthik Rajkumar, Rimpi Khurana, Edmond W Box, Yuguang Ban, Oliver G McDonald, Jashodeep Datta, Kathleen E DelGiorno, Nipun B Merchant, Sangeeta Goswami, Austin R Dosch, Nagaraj S Nagathihalli

**Affiliations:** Department of Surgery, University of Miami Miller School of Medicine, Miami, Florida, USA; Department of Public Health Sciences, University of Miami Miller School of Medicine, Miami, Florida, USA; Department of Pathology, University of Miami Miller School of Medicine, Miami, Florida, USA; Sylvester Comprehensive Cancer Center, University of Miami, Miami, Florida, USA; Sylvester Pancreatic Cancer Research Institute, University of Miami, Miami, Florida, USA; Department of Cell and Developmental Biology, Vanderbilt Ingram Cancer Center, Vanderbilt University, Nashville, TN, USA; Department of Genitourinary Medical Oncology and Immunology, The University of Texas MD Anderson Cancer Center, Houston, TX, USA

## Abstract

Pancreatic ductal adenocarcinoma (PDAC) remains a challenging disease in need of improved treatments. Cyclic adenosine monophosphate response element binding protein 1 (CREB) is an emerging therapeutic target whose oncogenic effects in PDAC have been largely attributed to a key molecular interplay between oncogenic *Kras^G12D/+^* (*Kras\**) and chronic inflammation driving irreversible acinar to ductal reprogramming. Here, we demonstrate that CREB activation fosters tumor associated macrophage (TAM) mediated immunosuppression and promotes PDAC growth in an aggressive *LSL-Kras^G12D/+^*;*Trp53^R172H/+^;Pdx1^Cre/+^*(*KPC*) genetically engineered mouse model. Selective deletion of CREB (*Creb^fl/fl^*) in *KPC*(*KPCC*^-/-^) mice attenuates primary disease burden. Unbiased transcriptomic analysis and validation using diverse molecular, genetic and pharmacological approaches *in vitro* and *in vivo* identify CREB-mediated transcriptional regulation of leukemia inhibitory factor (*Lif*) as one of the potential mediators of tumor cell-macrophage crosstalk promoting a pro-tumor polarization of TAMs, thereby attenuating the infiltration of effector T cells. Mechanistically, cancer cell derived LIF facilitates an immunosuppressive, pro-tumorigenic state. Importantly, pharmacological targeting of the CREB-LIF signaling axis between cancer cells and macrophages, using a CREB-specific inhibitor (CREBi), significantly suppresses tumor growth and sensitizes PDAC to immunotherapy, highlighting the therapeutic potential of this treatment combination to improve outcomes in this aggressive disease.

## Introduction

Pancreatic ductal adenocarcinoma (PDAC) is an aggressive solid cancer with a high mortality rate and is projected to become the second leading cause of cancer-related deaths by 2040 with a dismal 5 year survival rate of only 13% (1–3). Even today, PDAC remains to be refractory to nearly all conventional therapeutic regimens, demonstrating an urgent, unmet clinical need to develop new treatment strategies (4, 5). Contributing to this high degree of intrinsic and acquired therapeutic resistance in PDAC is a complex genetic landscape of transformed tumor cells, stromal desmoplasia, and a characteristic immunosuppressive tumor immune microenvironment (TME) established by immunosuppressive myeloid cell subsets and FOXP3^+^ T regulatory cells (6–8). As a result of this immunosuppressive phenotype, immune checkpoint-blockade (ICB) against programmed cell death protein-1(PD-1) or cytotoxic T-lymphocyte associated protein-4 (CTLA-4) has shown limited success in patients with PDAC (9).

Previous work have identified a role for specific driver mutations within the cancer cells, including *KRAS* and *TP53*, in activation of downstream effector signaling cascades that not only promote intrinsic tumor cell growth but also profoundly shape the diverse ecosystem of the TME (7, 10, 11). Despite growing recognition of the crosstalk between cancer cells, adjacent stromal and immune cells within the TME, the mechanisms by which this crosstalk drives PDAC remain largely unexplored. Therefore, identifying the key paracrine mediators that govern the composition of the TME holds immense potential in elucidating novel therapeutic targets and strategies to improve outcomes in PDAC by attenuating tumor progression and enhancing therapeutic efficacy of ICB immunotherapies.

Cyclic AMP response element-binding protein 1 (CREB) is a transcription factor known to mediate calcium, cytokine, and cellular stress response signaling (12), is activated by a multitude of upstream kinases including AKT, protein kinase A (PKA) and Mitogen- and Stress-activated protein Kinase (MSK1/2) (13–15). Once activated through phosphorylation at Ser-133, CREB binds to its coactivator, the CREB-binding protein (CBP), which enables recruitment of additional transcriptional machinery to fuel tumor growth (16). Notably, our recently published work has established a role for epithelial-cell intrinsic CREB activation as a key driver of PDAC initiation and progression with oncogenic *Kras* by coordinating a fibroinflammatory program within the pancreas in setting of chronic inflammation (17). CREB has also emerged as a vital transcription factor promoting high risk metastatic spread when concurrent with *TP53* alterations (11). However, the complexities of CREB signaling and its influence on the distinct stromal and immune components of the TME remains understudied.

Our present findings address this important knowledge gap by defining an epithelial cell intrinsic role for CREB in PDAC in driving immunosuppression and resistance to ICB. Herein, we comprehensively interrogate the role of CREB in promoting PDAC aggressiveness by integrating human pancreatic cancer multiomics datasets, preclinical genetically engineered mice models (GEMM) of *Creb* deletion (*Creb^fl/fl^*) in *LSL-Kras ^G12D/+^;Trp53^R172H/+^;Pdx1^Cre/+^*(*KPC*) mice, single cell RNA sequencing (scRNA-seq) analysis, CREB deleted murine orthotopic tumors, and pharmacological inhibition of CREB using the small molecule inhibitor 666-15. Our findings demonstrate that tumor cell intrinsic CREB contributes to macrophage mediated immunosuppression, in part, by regulating leukemia inhibitory factor (LIF) expression and signaling through its cognate receptor (LIFR) immunosuppressive macrophages. Importantly, we demonstrate that pharmacological inhibition of CREB using 666-15 remodels the TME to effectively reverse this immunosuppressive phenotype. By attenuating stromal fibroinflammatory responses and immunosuppressive myeloid-driven immune reprogramming, therapeutic targeting of CREB reverses T cell dysfunction and sensitizes otherwise immunorefractory preclinical PDAC models to immunotherapy, providing a sound rationale for this combination therapy as a novel treatment strategy to improve outcomes in PDAC patients.

## Results

### Cancer-cell autonomous CREB activation drives pancreatic tumorigenesis while genetic ablation attenuates disease burden in the KPC GEMM

We have previously shown that CREB activation in acinar cells promotes acinar to ductal metaplasia (ADM), thereby facilitating PDAC initiation and progression (17). To further establish its significance in human pancreatic cancer, we first conducted immunohistochemistry (IHC) analysis using a tissue microarray (TMA) representing a spectrum of pancreatic diseases (Figure 1*A*). Among the pancreatic tumor tissue cores analyzed, 50% (39/78) showed increased phosphorylated CREB (pCREB, Ser-133) activation localized within the tumor ductal compartment, whereas only 27% (19/70) of adjacent normal pancreatic tissue cores displayed increased pCREB expression (Figure 1*A*). Additionally, our analysis of publicly available human PDAC (PAAD) data from The Cancer Genome Atlas (TCGA) and GTEx datasets revealed overexpression of *CREB* mRNA in PDAC samples, (in both classical and basal subtypes) as compared to normal pancreatic tissues in healthy individuals (Figure 1*B*). Elevated *CREB* mRNA levels within these patient cohorts also correlated with poor progression-free survival underlying its clinical relevance in this malignancy (Figure 1*C*). In addition to TCGA PDAC datasets, we interrogated the Human Tumor Atlas Network (HTAN) dataset derived from treatment naïve PDAC patients (N=7). Using differentially expressed gene signatures, we attributed clusters to their putative identities, specifically tumor cell clusters were identified by *EPCAM* (epithelial), *KRT18*, and *KRT19* (ductal markers) expression (Supplementary Figure S1*A).* Uniform Manifold Approximation and Projection (UMAP) was used to display different intratumoral cellular compartments in these patients. Although *CREB* mRNA transcripts are localized to distinct subcellular compartments, its expression was predominantly enriched in tumor epithelial cells (Supplementary Figure S1*A*). Next, we delineated the biological significance of CREB expression in preclinical mouse models of PDAC. For this, we utilized pancreas tissues from age matched *Ptf1a^Cre/+^; LSL-Kras^G12D/+^* (*KC*) and *LSL-Kras^G12D/+;^ Trp53^R172H/+;^ Pdx1^Cre/+^*(*KPC*) mice (Figure 1*D*). These GEMMs encompass genomic alterations and capture the diverse spectrum of human PDAC progression, ranging from pancreatic intraepithelial neoplasia (PanIN) to invasive cancer, as previously described (18). *Pdx1-Cre* (*PC*) mice lacking oncogenic alleles were used as non-neoplastic controls (Figure 1*D*). Histological assessment using H&E, CK-19, and alcian blue staining revealed a markedly increased tumor burden in *KC* and *KPC* pancreatic tissues compared to non-tumor bearing (*PC*) pancreata, with the greatest extent of neoplastic transformation observed in *KPC* mice harboring both mutant *Kras* and *Trp53* co-alterations (Figure 1*D*). Interestingly, IHC analysis demonstrated an increase in the expression (activation) levels of phosphorylated CREB (pCREB, Ser133) within the pancreatic tumor ductal compartment of both *KC* and *KPC* mice, whereas minimal pCREB positivity was observed in the pancreata of control mice (Figure 1*D*). Additionally, RNA fluorescence in situ hybridization (RNA-FISH) employing a *Creb* mRNA probe set with protein immunofluorescence (IF) for CK-19, revealed the co-localization of *Creb* mRNA transcripts with CK-19 ductal cells in the pancreatic tumor sections of *KPC* GEMM mice (Supplementary Figure S1*B)*. Collectively, these findings provide compelling evidence for cancer cell intrinsic CREB activation in both human PDAC and preclinical GEMMs.

**Figure 1.**
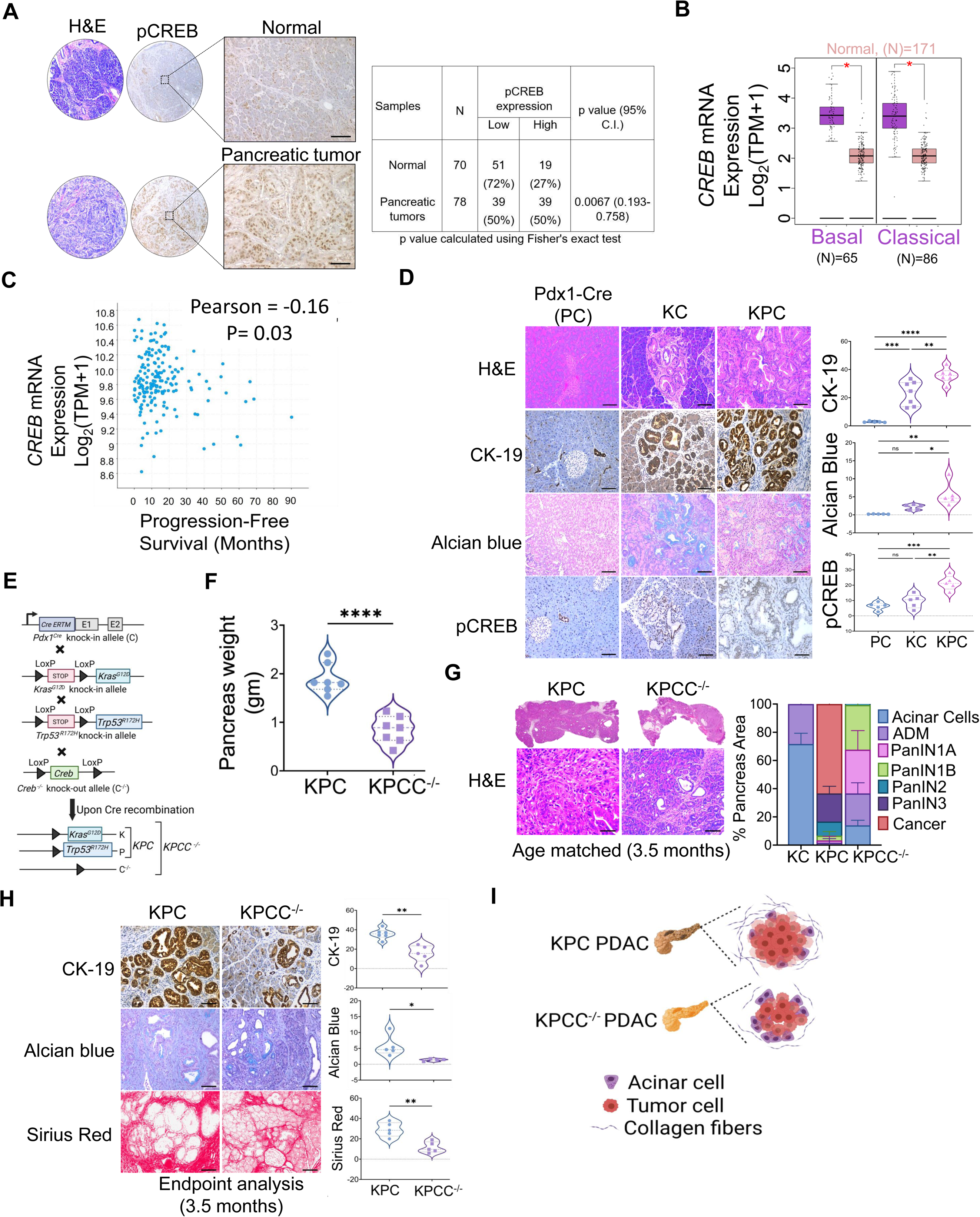
CREB is highly expressed in human and murine PDAC, and its genetic ablation attenuates tumor progression in *Kras^G12D/+^ Trp53^R172H/+;^Pdx1^Cre/+^*(*KPC*) genetically engineered mouse model (GEMM). **(A)** Representative H&E-stained pancreas sections and immunohistochemical (IHC) imaging analysis demonstrating pCREB (Ser-133) expression in the human tissue microarray of pancreatic tissue sections in (normal) controls and tumors *(left).* Quantitative analysis reveals heightened pCREB expression within the ductal epithelium of tumor samples compared to normal pancreas *(right)*. Two-sided Fisher’s exact test was performed. **(B)** mRNA levels of *CREB* in normal pancreatic tissue (Normal; n=171) vs. basal (n=65) and classic (n=86) subtype of PDAC patients (Tumor; n=151) from GTEx and PAAD TCGA dataset. **(C)** Pearson correlation plot depicting a significant association between high *CREB* mRNA expression and poor progression free survival in TCGA PDAC datasets (Tumor; n=179). **(D)** Representative H&E-stained pancreas sections and IHC imaging analysis along with quantification demonstrating CK-19, Alcian blue and pCREB expression in the pancreatic tissue sections of normal, and age matched tumor bearing *Ptf1a^Cre/+^; LSL-Kras^G12D/+^ (KC)* and *KPC* GEMMs of PDAC with n=5-7 mice per group. **(E)** Breeding strategy for generation of pancreas specific *Creb* deficient *KPC* (*Pdx1^Cre/+^; LSL-Kras^G12D/+^*; *Trp53^R172H/+^; Creb^fl/fl^*or *KPC*C^-/-^) **(F)** and comparative measurement of pancreas tumor weight in 3.5-month-old *Creb* wild type (*KPC*) and *KPC*C^-/-^ mice (n=7 mice per group). **(G)** Representative H&E images of the pancreata harvested from *KPC* and *KPCC^-/-^* mice along with comparative assessment using H&E-based histology to examine the entire pancreas, illustrating the presence of acinar cells, ADMs, PanINs, and cancerous regions in 3.5 months old *KPC* and *KPCC^-/-^* mice (n=3 mice per group). **(H)** Representative photomicrographs of the mouse pancreas sections imaging along with corresponding quantification demonstrating CK-19, Alcian blue and Sirius red staining in age matched tumor bearing *KPC and KPCC ^-/-^* GEMMs of PDAC with n=5-6 mice per group. (**I)** Schematic depicting pancreas specific deletion of *Creb* in *KPCC^-/-^* spontaneous mouse model attenuates pancreatic tumor progression and reduces fibrosis. Individual data points with mean ± SEM are shown and compared by two-tailed unpaired t test for two group comparison and one-way ANOVA for multiple comparisons; *p<0.050; **p<0.010; ***p<0.001; ****p<0.0001; ^ns^ non-significant (p>0.05). Scale bar, 50μm.

To further investigate the effects of CREB deletion on PDAC progression in the well-established *KPC* GEMM, we generated mice harboring a transgene that enables the expression of Cre recombinase under the control of the *Pdx-1* promoter (*Pdx1-Cre*) to knock in the mutant *Kras^G12D^* (*LSL-Kras^G12D/^*^+^) and *Trp53^R172H/+^* alleles to the endogenous loci in the murine pancreas (18). We then generated these *KPC* mice with homozygous loss of *Creb ^fl/fl^ (Creb*^−/−^) in the murine pancreas (Figure 1*E*). The two experimental groups: LSL-*Kras^G12D/+^;Trp53^R172H/+^;Pdx1^Cre/+^;Creb^+/+^*and *LSL-Kras^G12D/+^;Trp53^R172H/+^;Pdx1^Cre/+^;Creb^fl/fl^*(subsequently will be referred to as *KPC* or *KPCC^-/-^*, respectively, throughout the text). Examination of the pancreata from *KPCC^-/-^*mice revealed absence of *Creb* mRNA as compared to the pancreatic tumor tissue from *Creb* wild-type *KPC* mice (Supplementary Figure S1*C)*. Notably, the tail tissue of the mice displayed intact *Creb* expression, confirming the tissue specificity of the *Pdx1-Cre* promoter (Supplementary Figure S1*C)*. Additionally, RNA-FISH analysis using a *Creb* mRNA probe set in conjunction with CK-19 demonstrated a loss of mRNA transcripts within the ducts, confirming *Creb* ablation in the pancreas of *KPCC^-/-^* mice as compared to *KPC* (Supplementary Figure S1*D)*.

Pancreatic tissues were collected from these two cohorts at 3.5 months of age for end point analysis to measure primary disease burden. Intriguingly, a marked reduction in pancreas tumor weight was observed in *KPCC^-/-^*as compared to *KPC* (Figure 1*F*). Next, on histological examination, pancreata from *KPC* mice displayed a higher prevalence of high grade PanIN lesions and invasive cancer as compared to *KPCC^-/-^* mice (Figure 1*G*). Analyses of age matched *KC* mice harboring *Kras^G12D/+^ (Kras*)* alone exhibited only a few ADM lesions within a large predominance of normal acinar architecture. Notably, *Creb*-deleted pancreata of 3.5-month-old *KPCC^-/-^*mice were devoid of high-grade lesions and displayed less PDAC disease burden. Additionally, detailed histological analysis of pancreatic tissue sections stained with CK-19 and alcian blue revealed microscopic changes indicative of diminished disease progression in *KPCC^-/-^* mice, relative to *KPC* mice (Figure 1*H*). Sirius red staining showed a marked decrease in the fibrotic content of pancreata of *KPCC^-/-^*compared to *KPC* mice (Figure 1*H*).

To investigate CREB-dependent spatial changes in the TME of *KPC* tumors, we performed NanoString GeoMx digital spatial profiler (DSP) in *KPC* vs *KPCC^-/-^* pancreata (Supplementary Figure S1*E)*. Use of two antibodies targeting immunosuppressive macrophages (F4/80) and tumor epithelial cells (PanCK), along with DAPI, resulted in compartmentalization of macrophage and epithelial (tumor)-enriched regions (Supplementary Figure S1*E)*. GeoMx DSP analysis identified Ki-67, phosphorylated c-Jun-N-terminal kinase (pJNK), V-domain Ig suppressor of T cell activation (Vista), and fibronectin as among the downregulated proteins in the pancreata of *KPCC^-/-^* compared to *KPC* mice (data combined from both cellular compartments). IHC validation of these findings further confirmed significant downregulation of the Ki67^+^ proliferative index in these tissues supporting the observed decrease in tumor burden with CREB deletion (Supplementary Figure S1*F)*. Taken together, these findings demonstrate the critical involvement of CREB in dictating primary disease burden and progression in the *KPC* GEMM of PDAC through the suppression of tumor cell growth and proliferation and a reduction in stromal fibrosis, suggesting a vital role for tumor cell-derived CREB in mediating these processes (Figure 1*I*).

### CREB orchestrates tumor macrophage crosstalk to promote an immunosuppressive microenvironment

Immunosuppression in PDAC is driven, in part, by sustained activation of cancer cell intrinsic oncogenic pathways that orchestrate complex autocrine and paracrine crosstalk with the TME (7, 19). To determine the role of tumor-cell derived CREB in mediating these processes, we first interrogated scRNA-seq datasets from treatment-naïve human pancreatic cancer patients (N=7 patients generated by the human tissue atlas network (HTAN) (Figure *2A-C*). Intercellular communication networks were quantitatively inferred using the CellChat algorithm within tumor cells subclusters with high or low *CREB* expression (Figure *2A*). CellChat analysis identified 15 different cell type clusters (Figure *2B*). To quantify outgoing cell communication from CREB high tumor cells to other cell types, we subset the interactions where CREB high tumor cells were the source and filtered for significance (p<0.050) (Figure *2C*). Our analysis identified statistically significant outgoing communication with a range of immune and stromal cell types, the strongest aggregated interaction based on total communication probability across significant ligand receptor pairs (p<0.050) was observed with B cells, endothelial cells, and monocytes/macrophages population (Figure *2C*). Though B cells showed the highest overall target, monocytes/macrophages remained among the top ranked targets supporting their relevance as a key recipient of CREB high tumor derived signals and aligning with their central role in modulating the tumor immune microenvironment.

**Figure 2.**
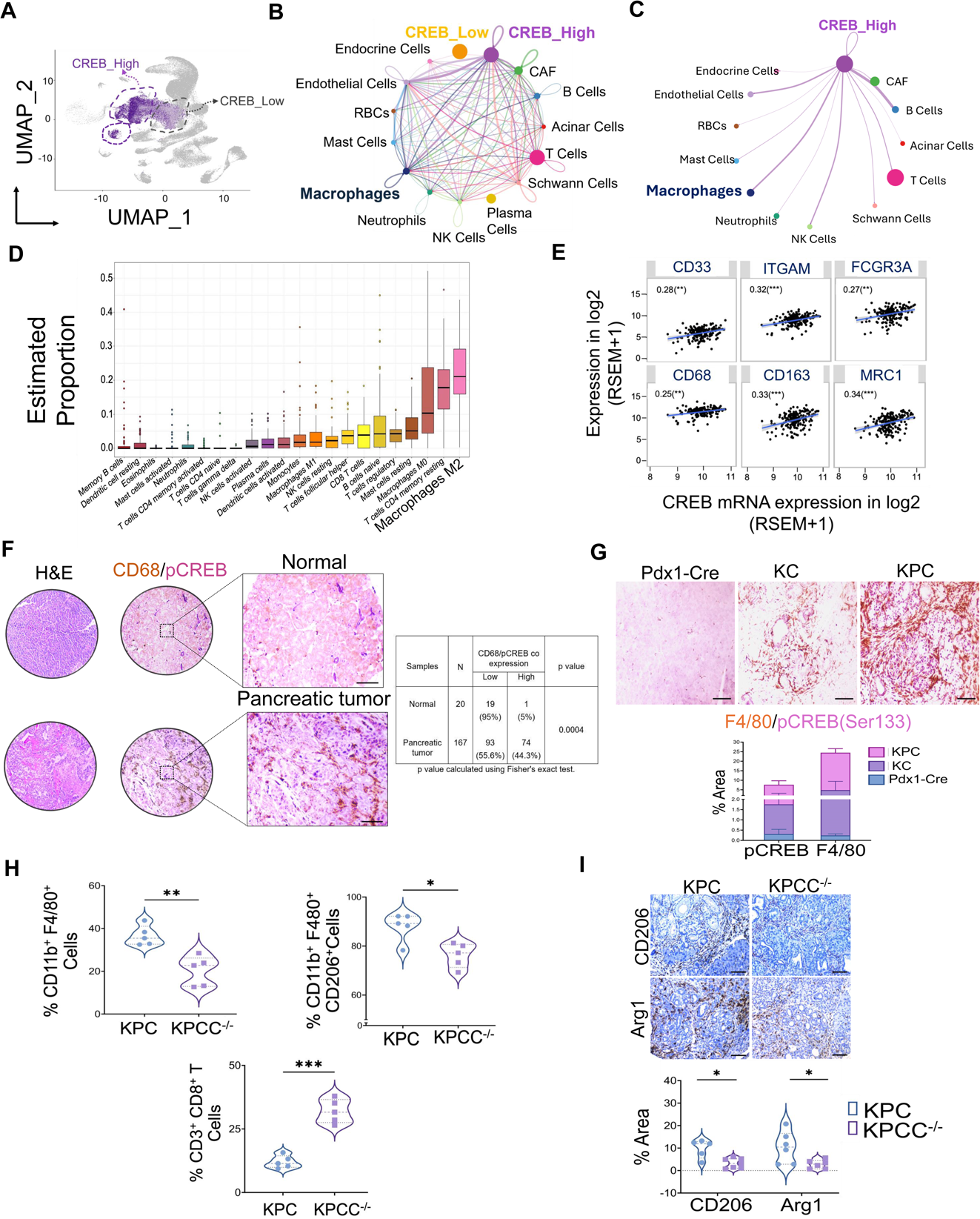
CREB orchestrates tumor–macrophage crosstalk and promotes an immunosuppressive microenvironment. **(A)** Visualization of sub-clustering in the human HTAN dataset based on CREB activity gene scores within tumor cell clusters. UMAP: Uniform Manifold Approximation and Projection. **(B-C)** Results of the Cell-Chat analysis for important ligand receptor pairs linking diverse cellular communications. **(D)** Immune cell deconvolution analysis of pancreatic cancer in TCGA PAAD cohort revealing estimated proportions of immune cellular composition in these patients. **(E)** Spearman correlation analysis revealing significant association of *CREB* mRNA expression with myeloid/macrophage markers in TCGA PAAD cohort. **(F)** Representative photomicrographs of H&E and dual color IHC staining with corresponding quantification for pCREB/CD68 (pan macrophage marker) of a human TMA depicting high co-expression of pCREB within the tumor cells, alongside CD68^+^ macrophage infiltration in pancreatic tumor tissue sections when compared to normal. A two-sided Fisher’s exact test was performed for statistical significance. **(G)** Representative photomicrographs of dual color IHC staining with corresponding quantification for pCREB/F4/80 (pan macrophage marker) expression within the pancreatic tissue sections of normal (*Pdx1-Cre*) and age matched tumor bearing *Ptf1a^Cre/+^; LSL-Kras^G12D/+^ (KC)* and *KPC* GEMMs of PDAC with n=4 mice per group. **(H)** Flow cytometric analysis demonstrates significant reduction in the percentage (%) of TAMs (CD11b^+^ F4/80^+^), and M2-like (CD11b^+^ F4/80^+^ CD206^+^) macrophages in the pancreatic tumors with *Creb* deletion in *KPCC ^-/-^* GEMM compared to wild type *KPC* mice with n=5 mice per group *(top)*. Flow cytometric analysis (%) of tumor infiltrating T cell adaptive immune compartment depicting frequency of CD8^+^ T cells (CD45^+^ CD3^+^ CD8^+^) in the pancreatic tumors with *Creb* deletion in *KPCC ^-/-^* GEMM compared to wild type *KPC* mice with n=5 mice per group *(bottom)*. **(I)** Representative photomicrographs of the mouse pancreas sections imaging along with corresponding quantification demonstrating staining of TAMs markers (Arg1, CD206) in age matched tumor bearing *KPC* and *KPCC^-/-^* GEMMs of PDAC with n=5-7 mice per group. Individual data points with mean ± SEM are shown and compared by two-tailed unpaired t test. *p<0.05; **p<0.01; ****p<0.0001; ^ns^ non-significant (p>0.05). Scale bar, 50μm.

Within the TME, immunosuppressive macrophages are the most abundant myeloid cell subset, and they are critical players in orchestrating crosstalk with cancer cells through molecular programs that govern innate immune subset trafficking and function (7, 20, 21). Performing CIBERSORT analysis on the TCGA PDAC dataset, we estimated the proportion of tumor infiltrating immune cell subsets and identified 22 subsets in these samples, with tumor associated macrophages (TAMs) and alternatively activated immunosuppressive macrophages emerging as the most prominent populations (Figure *2D*). To further establish the significance of CREB expression and its relationship with TAMs, we next examined the correlation between *CREB* mRNA expression levels and gene signatures of innate myeloid/macrophages in human pancreatic cancer patient samples (cohort PDAC-TCGA, n=178) (Figure *2E*). Our analysis revealed a significant positive correlation between *CREB* expression and gene signatures associated with immunosuppressive myeloid/macrophage infiltrated immune contexture comprised of *CD33, ITGAM, FCGR3A, CD68, CD163,* and *MRC1*. To further extend these findings in pancreatic cancer, we performed dual-color IHC using a pancreas TMA with a total of 96 cases/192 cores (Figure *2F*). Pancreatic cancer tissues displayed a significant increase in CD68^+^ macrophage infiltration compared to adjacent normal pancreas. Additionally, this immunosuppressive myeloid/macrophage-rich TME was associated with ductal cell-specific hyperactivation of pCREB in tumor sections in these patients. Similarly, analysis of pancreas tissue sections from GEMMs (*KC* and *KPC* mice) revealed an increase in activation of pCREB with heightened TAMs (F4/80^+^) infiltration, particularly in *KPC* with increased tumor burden when compared to either *KC* or non-tumor-bearing control mouse pancreas (Figure *2G*).

Furthermore, flow cytometric immunophenotyping analysis of mouse pancreatic tumors harvested from *KPC* and *KPCC^-/-^* mice displayed a significant decrease in the frequency of both total macrophages (CD11b^+^ F4/80^+^ TAMs) and alternatively activated immunosuppressive macrophages with an M2-like phenotype (CD11b^+^ F4/80^+^ CD206^+^) in *KPCC^-/-^* tumors compared to *KPC* (Figure *2H*). Reduced myeloid/macrophage remodeling was associated with a significant increase in the frequency of CD3^+^ CD8^+^ T-cell populations in *KPCC^-/-^* pancreata (Figure 2*H*). Further validation using IHC on harvested pancreatic tumor sections from the two groups (*KPC* vs. *KPCC^-/-^* GEMM) (Figure 2*I*) confirmed observations made with flow cytometry demonstrating a decrease in the expression of TAMs-associated immunosuppressive signatures, including ARG1 and CD206, within the pancreata of *KPCC^-/-^* compared to *KPC*. Taken together these findings provide scientific evidence supporting the critical role of CREB in shaping the immunosuppressive microenvironment associated with TAMs in PDAC.

### Genetic silencing of tumor cell intrinsic CREB overcomes macrophage mediated immunosuppression and enhances T cell infiltration in the orthotopic KPC mice model

To elucidate the dynamics of CREB mediated tumor cell macrophage crosstalk, within the TME, we utilized *KPC* tumor cells in which *Creb* has been genetically silenced using CRISPR-*Cas9* based genome editing (Supplementary Figure S2*A*). Targeted silencing of CREB expression in a single clonally expanded cell population of *KPC* mouse tumor cells was validated using Western blot (Supplementary Figure S2*A*). A syngeneic orthotopic model of *KPC* was established by implanting either *KPC*-*Creb* wild type (CREB^WT^) or *Creb* knockout (CREB^KO^) tumor cells in the pancreata of C57BL/6 mice (Figure 3*A*). ScRNA-seq was then performed on these tumors 4 weeks after initial implantation, to capture cellular and transcriptional heterogeneity using the 10x Genomics platform (Figure 3*A*). By utilizing differentially expressed gene signatures, we successfully attributed clusters to their putative 9 cellular identities, with macrophage and T cell clusters being designated by specific gene expression patterns (Supplementary Figure S2*B)*. UMAP was used to visualize distinct intratumoral cellular compartments within the pancreas (Figure 3*B*).

**Figure 3.**
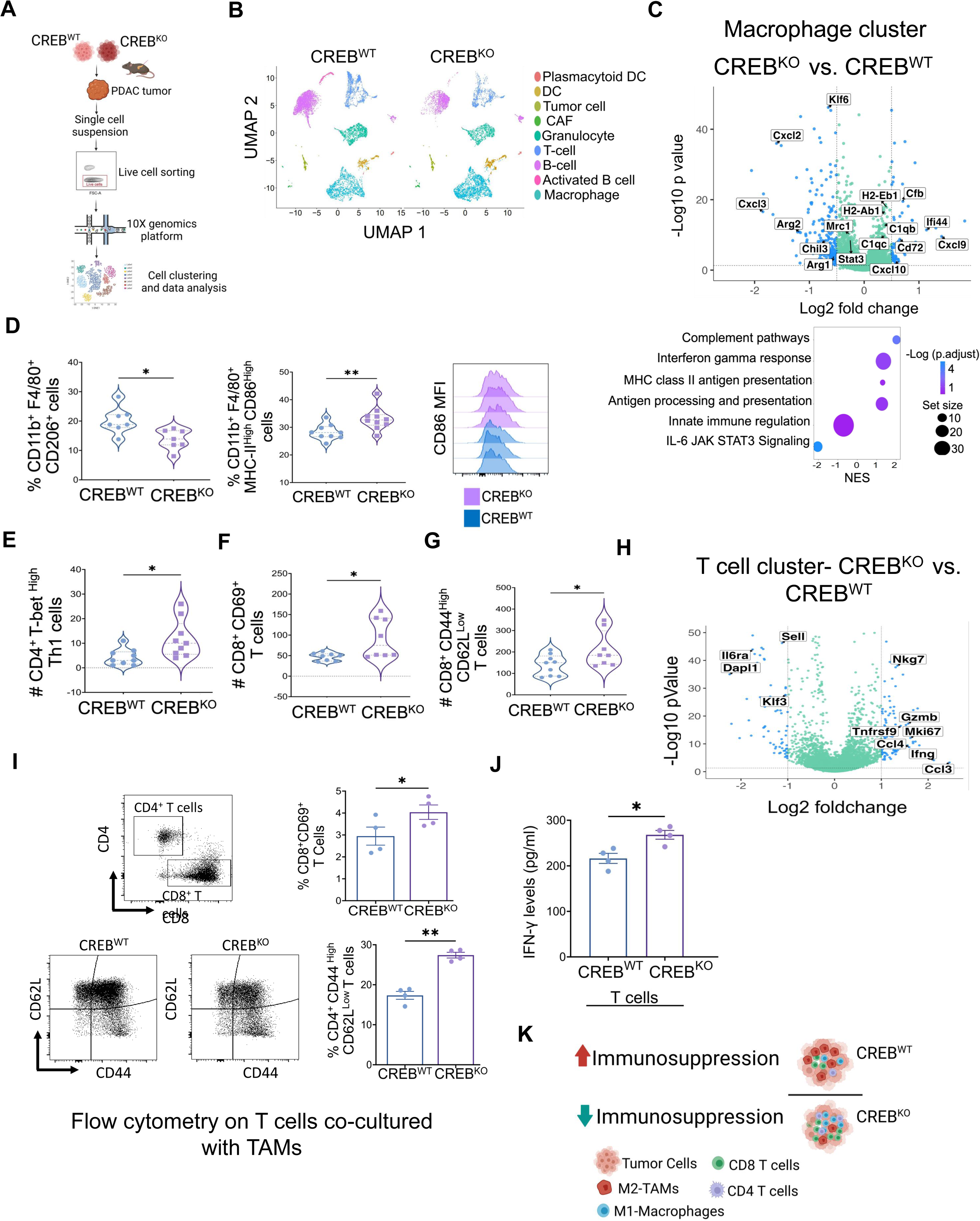
Genomic ablation of CREB modulates TAMs and facilitates intratumoral T cell infiltration in an orthotopic KPC mouse model of PDAC. **(A)** Experimental schematic of single cell RNA sequencing (scRNA-seq) workflow conducted on CREB^WT^ and CREB^KO^ orthotopic mouse PDAC tumors using 10X genomics platform. **(B)** UMAP projection plot displaying cell clusters of multiple cell types present within the TME of PDAC. **(C)** Volcano plot illustrating differentially expressed genes (DEG) *(top)* and bubble plot showing gene set enrichment analysis (GSEA) in the macrophage cell subcluster of CREB^KO^ vs. CREB^WT^ tumors in KPC mice *(bottom).* **(D)** Flow cytometric analysis demonstrates significant reduction in the percentage (%) of M2 like TAMs (CD11b^+^ F4/80^+^ CD206^+^), with a parallel increase of antigen presenting M1-like (CD11b^+^F4/80^+^MHC-II^High^CD86^High^) macrophages in the pancreatic tumors with CREB deletion as compared to wild type mice with (n=7-10) mice per group *(left)*. Representative mean fluorescence intensity (MFI) histogram showing high CD86 expression on macrophages of CREB deleted (CREB^KO^) pancreatic tumors *(right)*. **(E-G)** Flow cytometric assessment of the tumor-infiltrating adaptive immune compartment depicting proportion of CD4^+^ Tbet^High^ cells (Th1), activated CD8^+^ (CD69^+^ CD8^+^) T cells and effector memory CD8^+^ (CD8^+^ CD44^High^ CD62L^Low^) T cells in CREB^WT^ and CREB^KO^ orthotopic mouse PDAC tumors with n=8-9 mice per group. **(H)** Volcano plot illustrating DEG in the T cell subcluster of CREB^KO^ vs. CREB^WT^ tumors in KPC mice. **(I)** Flow cytometric analysis showing a significant increase in activation of CD4^+^ and CD8^+^ T cell phenotypes in an *ex vivo* coculture system with TAMs isolated from CREB^WT^ or CREB^KO^ KPC orthotopic tumors with pan T cells (n=4 per group). **(J)** Quantification of IFN-γ secretion by ELISA in supernatants from the same co-culture systems with T cells and CREB^WT^ or CREB^KO^ TAMs after 48hr of co culture (n=4 per group). **(K)** Schematic depicting tumor cell intrinsic deletion of CREB in KPC in an orthotopic mouse model attenuates immunosuppression. Individual data points with mean ± SEM are shown and compared by two-tailed unpaired t test. *p<0.05; **p<0.01; ^ns^ non-significant (p>0.05). Scale bar, 50μm.

Notably, compared with the CREB^WT^ tumors, tumor cell-intrinsic deletion of CREB resulted in substantial reprogramming of the macrophage transcriptome. Differentially expressed gene analysis of the macrophage transcriptome of CREB^KO^ tumors revealed significant downregulation of gene signatures linked to an immunosuppressive and potentially tumor promoting phenotype (*Arg1, Chil3, Cxcl2, Cxcl3, Mrc1*), with a concomitant positive enrichment of genes highlighting antigen presentation (*H2-Ab1*), the IFN-γ response (*Cd44*), and enhanced activation and recruitment of T cells (*Cxcl9, Cxcl10*) (Figure 3*C*, *top*), indicating a switch to a more anti-tumor phenotype. We then performed pathway enrichment analysis between CREB^KO^ vs. CREB^WT^ within the macrophage cluster. This analysis revealed positive enrichment of pathways related to Class II MHC antigen presentation and response to IFN-γ alongside significant downregulation of IL-6–JAK–STAT3 signaling, further indicative of a shift towards a more immunostimulatory and anti-tumorigenic macrophage phenotype in the absence of tumor cell-intrinsic CREB (Figure 3*C*, *bottom*).

To further validate our scRNA-seq findings and investigate whether cancer cell intrinsic CREB promotes TAM polarization towards an immunosuppressive phenotype, we performed multiplex immunophenotyping analysis looking into macrophage populations in CREB^WT^ vs CREB^KO^ orthotopic pancreatic tumors (Figure 3*D*). Our analysis revealed a significant reduction in the tumor infiltrating alternatively activated TAM population (CD11b^+^F4/80^+^CD206^+^) and an increase in antigen presenting classically activated macrophages with an M1-like phenotype (CD11b^+^F4/80^+^MHC-II^High^CD86^High^) in CREB^KO^ as compared to CREB^WT^ (Figure 3*D*). Additional validation of these findings using qPCR-based mRNA assessment in bulk tumors revealed significant reduction in the expression of M2 related signature marker expression (CD206 and Arg1) in CREB deleted tumors compared with wild type (Supplementary Figure S2*C*). Notably, we also observed a significant increase in the mRNA transcript profile of the M1 marker (inducible nitric oxide synthase *(iNOS/Nos2)*) in CREB deleted tumors as compared to wild type (Supplementary Figure S2*D).* Additionally, in alignment with the increased transcriptional signatures of activated T cell recruiting chemokines observed in CREB^KO^ tumors, qPCR analysis of bulk tumor tissue further demonstrated elevated mRNA expression of *Cxcl9* and *Cxcl10* in CREB deleted bulk tumor profiles as well, suggesting a possible link between CREB deletion and a heightened T cell chemotactic gradient within the TME (Supplementary Figure S2*E)*.

Next, we aimed to determine whether the alterations associated with CREB silencing affected the T cell-based adaptive immune response in our *KPC* orthotopic PDAC model (Figure 3*E-H*). We observed a significant increase in intratumoral CD4^+^ Th1 cells (CD4^+^Tbet^High^) (Figure 3*E*), and activated CD8^+^ (CD8^+^CD69^+^, Figure 3*F*) and effector memory CD8^+^ (CD44^+^ CD62L^-^) (Figure 3*G*) T cells within the TME of CREB^KO^ tumors as compared to wild type. Furthermore, transcriptional profiling of the T cell cluster from scRNA-seq of CREB^KO^ tumors revealed downregulation of genes expressed in naive populations and associated with quiescence (*Klf3, Il6ra, Dapi1)* with enrichment of gene signatures highlighting activated, cytotoxic, and effector memory T cell states (*Gzmb*, *Ifng, Ccl4, Ccl3),* (Figure 3*H*).

To determine whether macrophage polarization, driven by tumor cell intrinsic CREB, directly impairs T cell activation and function, we isolated F4/80^+^ TAMs from CREB^WT^ and CREB^KO^ tumors (n=4 mice in both group) and co-cultured these populations *ex-vivo* for 48 hours with anti-CD3/CD28 stimulated T cells isolated from non-tumor bearing C57BL/6 mice (Supplementary Figure S2*F* and Figure 3*I*). Flow cytometric analysis showed that CREB^KO^ TAMs promoted activation of (CD45^+^ CD11b^-^ TCRβ^+^) CD8^+^ CD69 ^+^ T cells along with increased frequency of CD4^+^ T cells with an effector memory phenotype (CD44^high^ CD62L^low^). Functionally, T cells cocultured with CREB^KO^ TAMs showed an enhanced cytotoxic profile (Figure 3*J*) as evidenced by a significant increase in IFN-γ secretion. Overall, these results suggest that silencing tumor cell intrinsic CREB effectively overcomes the immunosuppressive TME by influencing macrophage polarization and promoting T cell infiltration and activation in PDAC GEMMs (Figure 3*K*).

### Tumor cell intrinsic CREB mediated transcriptional regulation of the leukemia inhibitory factor

To elucidate the mechanism(s) underlying CREB-mediated TAM modulation, we conducted RNA sequencing (RNA-seq) analysis in CREB^WT^ and CREB^KO^ *KPC* tumor cells (Figure 4*A-B*). Gene set enrichment analysis in CREB^KO^ cells revealed a significant negative enrichment of oncogenic signaling pathways involved in tumor progression and aggressiveness, including *(Kras, Tp53*, *TNF,* and EMT signaling) (Figure 4*A*) as well as downregulation of leukemia inhibitory factor (*Lif*) (Figure 4*B*), a secreted factor implicated in TAM-mediated immunosuppression through activation of JAK/STAT3 signaling by binding to the cognate gp130/LIFR complex at the target cell surface (22, 23). Indeed, GSEA analysis of the human HTAN PDAC dataset identified this signaling node within the macrophage compartment (Figure 4C). To validate a role for LIFR signaling in immunosuppressive macrophages, we employed mouse RAW macrophages (264.7) and stimulated with recombinant LIF (rLIF), which led to activation of pSTAT3 (Tyr705); using a small molecule inhibitor and antagonist of LIF receptor (EC359) abrogated this activation (Figure 4*C*-*D*). Collectively, these results suggest that tumor cells-specific CREB drives immunosuppression through LIF-mediated paracrine communication which activates JAK-STAT3 signaling in immunosuppressive macrophages.

**Figure 4.**
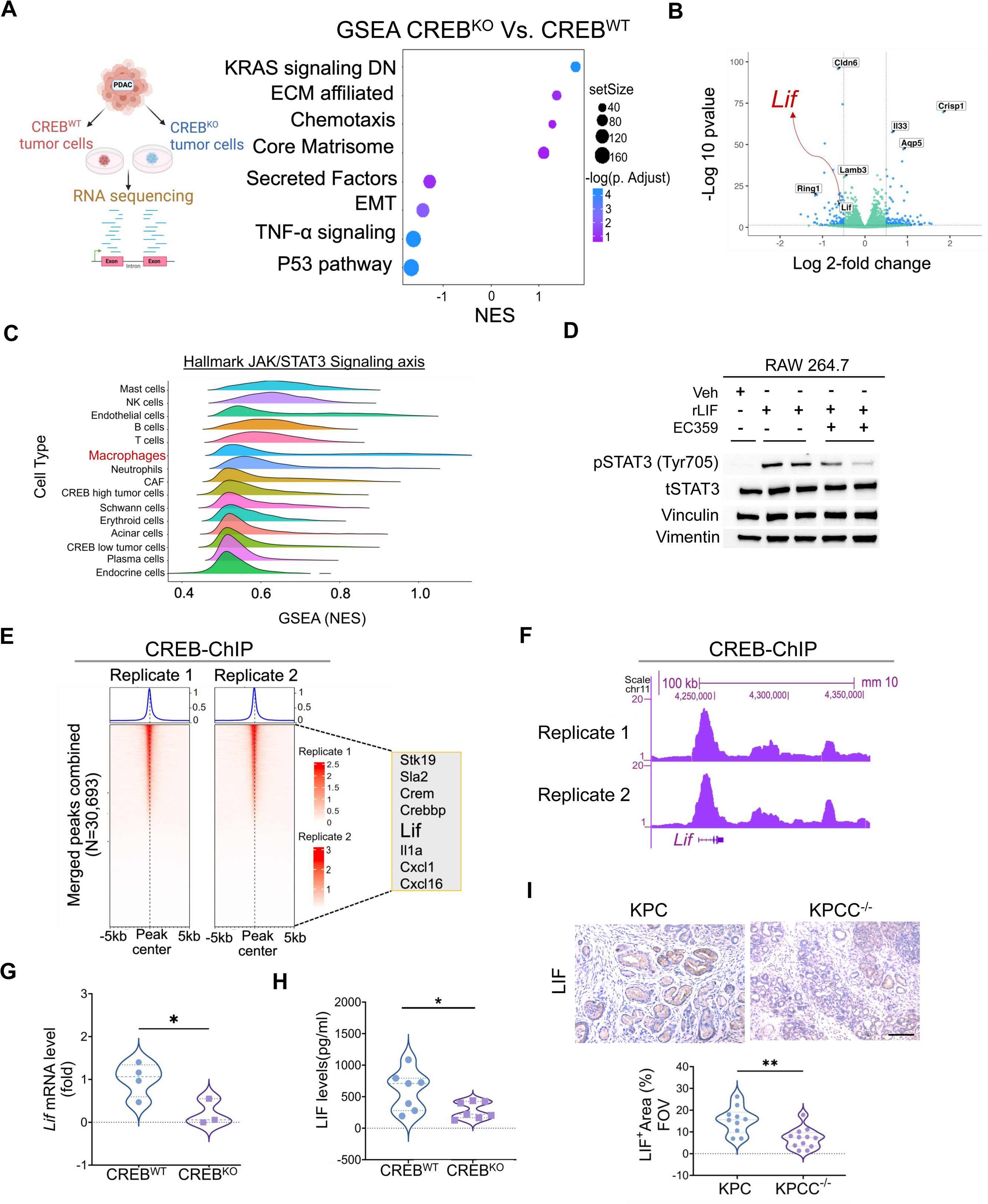
Cancer cell intrinsic CREB transcriptionally regulates LIF expression, driving macrophage-modulatory signaling in PDAC. **(A)** Experimental schematic demonstrating RNA sequencing (RNA-seq) analysis conducted in KPC mouse CREB^WT^ and CREB^KO^ tumor cells. **(B)** Bubble plot visualizing differentially regulated pathways (using molecular signature databases) in CREB^KO^ transcriptomics compared with CREB^WT^ tumor cells (n=3). **(C)** Ridge plot showing normalized enrichment scores for the hallmark JAK–STAT3 signaling pathway across different cellular constituents in the human HTAN pancreatic cancer dataset from treatment-naïve patients. **(D)** Western blot image depicting pSTAT3 activation, along with tSTAT3, vinculin and vimentin in mouse RAW macrophage 264.7 cell line treated with either recombinant (r) LIF alone (1 µg/mL) or in combination with EC359 (LIFR blockade, 0.2µM). **(E)** Chromatin Immunoprecipitation sequencing (ChIP-seq) peak signals visualized as heat maps which depict CREB binding sites across genomic regions in KPC mouse pancreatic tumor cells. The adjacent call out boxes with the heat maps showing essential genes regulated via CREB as its downstream mediators. **(F)** Integrative Genome Viewer (IGV) plot visualizing occupancy of CREB binding peaks in ChIP-seq data at the site of mouse *Lif* promoter gene regulatory sequences. **(G-H)** Violin dot plots showing downregulation of LIF expression via qPCR and ELISA in bulk tumor lysates of CREB^KO^ KPC orthotopic PDAC tumors as compared to CREB^WT^ with n=3-7 mice per group. **(I)** Representative photomicrographs of whole pancreas along with its corresponding quantification depicting significantly reduced LIF expression via IHC in age matched *KPC*C^-/-^ as compared to *KPC* mice. Individual data points with mean ± SEM are shown and compared by two-tailed unpaired t test. *p<0.05; **p<0.01; ^ns^ non-significant (p>0.05).

To explore the regulatory role of CREB on LIF expression, we utilized the CREB^WT^ and CREB^KO^ tumor cells. Both *Lif* gene expression and secreted protein LIF levels were significantly reduced in CREB^KO^ tumor cells, compared to CREB^WT^ (Supplementary Figure S3*A* and *B*). Additionally, *KPC* tumor cells transduced with a *Creb* lentiviral vector exhibited a significant increase in LIF secretion in the conditioned media of CREB overexpressed cells (CREB^OE^) cells as compared to cells transduced with non-targeting control vector (CREB^NTV^) (Supplementary Figure S3*C*).

Analysis of the human PDAC TCGA and GTEx datasets revealed overexpression of *LIF* mRNA in pancreatic tumor samples as compared to normal tissues (Supplementary Figure S3*D*), thereby suggesting a putative role for LIF in this tumor type. To determine whether CREB plays a direct role in regulating *Lif* expression, we performed chromatin immunoprecipitation sequencing (ChIP-seq) in *KPC* tumor cells (Figure 4*E-F*). ChIP-seq analysis of CREB enriched genomic regions revealed many putative CREB targets involved in immunomodulation, tumor cell proliferation, and migration (*Il1a, Cxcl1, Cxcl16, Stk19, Sla2, Crem, Crebbp*, (Figure 4*E*). Interestingly, UCSC genome browser ChIP-seq visualization showed a distinct ChIP-seq signal for CREB at *Lif* gene region, providing evidence for CREB binding on *Lif* (Figure 4*F*). We also noted a significant reduction in *Lif* mRNA expression and secretion in bulk tumor lysates prepared from harvested from CREB^KO^ tumors, in comparison to CREB^WT^ (Figure 4*G*-*H*). IHC-based histological profiling on the pancreata of *KPC* mice revealed elevated LIF expression within ADM/PanIN lesions, while the pancreata of CREB deleted *KPCC^-/-^*mice displayed significantly attenuated expression of LIF, providing additional confirmation of CREB mediated regulation of this cytokine (Figure 4*I*).

To assess the clinical significance of CREB-LIF co-expression, we analyzed publicly available TCGA PDAC datasets, which demonstrated a significant positive correlation between *LIF* and *CREB* mRNA expression with a Pearson correlation coefficient of R=0.76, (Supplementary Figure S3*E*). Additionally, patients with elevated expression of both *LIF* and *CREB* mRNA, based on combined gene signature, displayed significantly reduced disease-free survival as compared to those with lower expression levels (p=0.032, HR=1.6) (Supplementary Figure S3*F*), highlighting that their combined activity may promote poor clinical prognosis in PDAC.

Additional validation of mRNA expression data from 35 human PDAC cell lines in the CCLE dataset via DepMap portal also revealed a significant positive correlation between *CREB* and *LIF* mRNA expression levels (R=0.36, p=0.031) (Supplementary Figure S3*G*), supporting a coordinated relationship. Furthermore, qPCR-based transcriptional profiling of six human pancreatic cancer cell lines and one non-cancerous immortalized pancreatic cell line as control (HPNE) (Supplementary Figure S3*H*) revealed broad expression of the *CREB* and *LIF* mRNA axis across pancreatic cancer cell lines. Collectively, these findings underscore the role of cancer cell autonomous CREB expression in regulating *Lif* and the clinical relevance of the *CREB-LIF* signaling axis in PDAC.

### CREB driven LIF expression from tumor cells orchestrates tumor-macrophage cross talk in PDAC

We first screened for LIFR expression in different cell populations within the TME. Flow cytometric analysis showed TAMs are the predominant cell population expressing LIFR, when compared to other populations present within the TME of orthotopic *KPC* tumors (Figure 5*A*). Next, we aimed to study the direct role of CREB-regulated LIF in driving TAM polarization.

**Figure 5.**
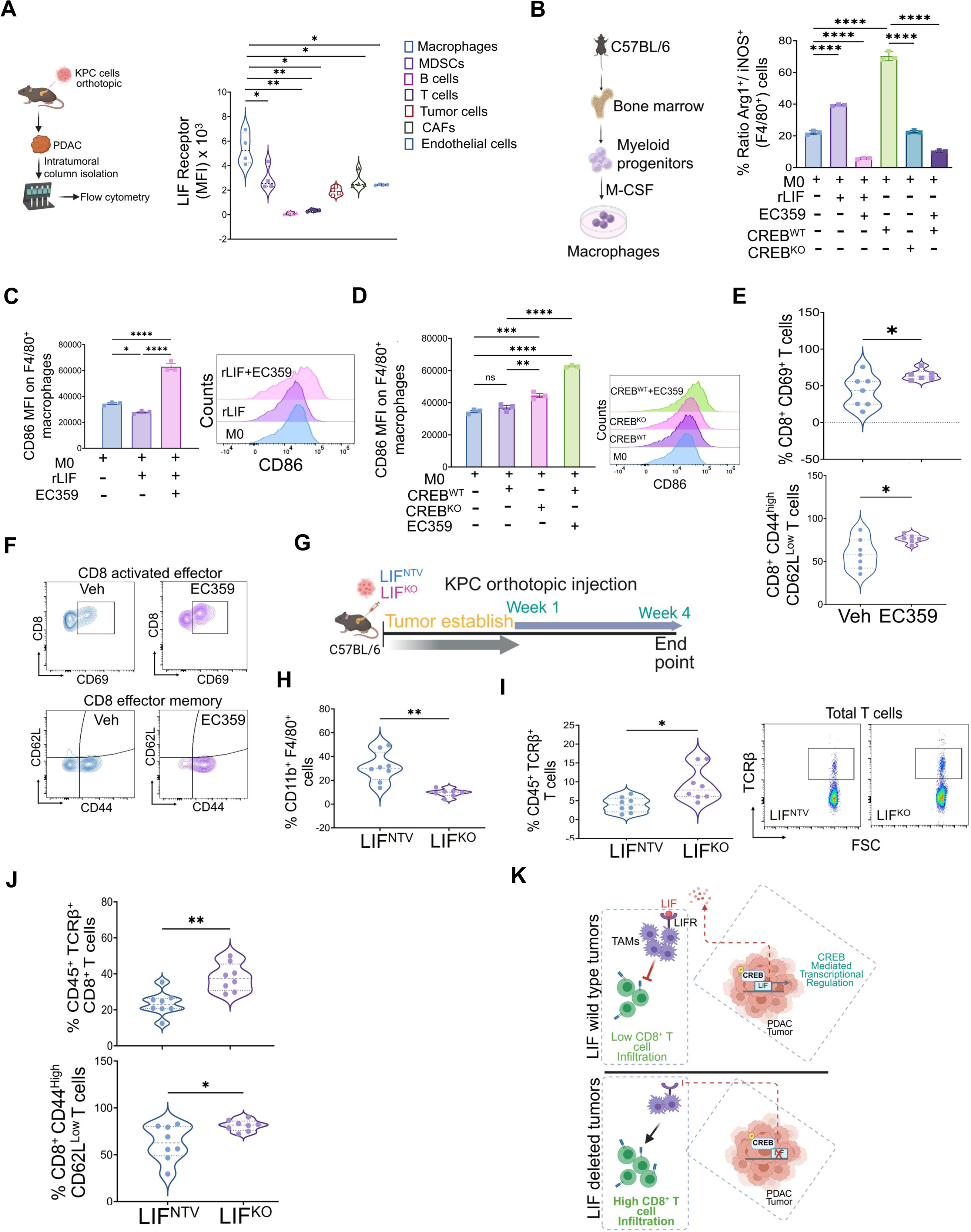
Cancer cell intrinsic CREB-LIF mediates tumor-macrophage cellular crosstalk in PDAC. **(A)** Experimental schematic demonstrating intratumoral isolation of different cellular constituents from KPC orthotopic tumors and subsequent flow cytometric based assessment *(left)*. Violin plots depicting the highest expression of LIF receptor (LIFR) on the F4/80^+^ macrophages as compared to other cell types within the TME (*right*). **(B)** Schematic showing differentiation of bone marrow derived myeloid progenitors into macrophages (BMDMs) stimulated by M-CSF in C57BL/6 mice *(left)*. Bar plot of flow cytometric analysis depicting the ratio of Arg1^+^/iNOS^+^ (F4/80^+^) cells in BMDMs under various experimental conditions (*right*). These include treatment with recombinant LIF (rLIF) either alone or in combination with LIFR blockade (EC359); conditioned media from KPC CREB^WT^ tumor cells either alone or in combination with EC359; conditioned media from CREB^KO^ cells. **(C)** Bar plot of flow cytometric analysis depicting mean fluorescence intensity (MFI) of CD86 expression on F4/80^+^ macrophages treated with recombinant LIF (rLIF) either alone or in combination with EC359 (*left*) along with representative MFI histogram plots (*right*). **(D)** Bar plot of flow cytometric analysis depicting MFI of CD86 expression on F4/80^+^ macrophages treated conditioned media from CREB^WT^ tumor cells either alone or in combination with EC359; conditioned media from CREB^KO^ cells, along with demonstration of representative MFI histogram plots (*right*). **(E)** Violin plot of flow cytometric analysis depicting a significant increase in the infiltration of activated CD8^+^ T cells with an effector memory phenotype following LIFR blockade (EC359) in an *in vivo* KPC orthotopic PDAC mouse model (n=6-7 mice per group). **(F)** Representative image of the contour plots depicting shift in the population of CD8^+^ CD69^+^ T cells and CD8^+^ CD44^high^ CD62L^low^ (effector memory) T cells in EC359 treated pancreatic tumors compared to vehicle. **(G)** Schematic illustrating orthotopic tumor cell implantation of KPC cells harboring wild type LIF expression (LIF^NTV^) or LIF^KO^ into the pancreas of C57BL/6 mice. **(H)** Flow cytometric analysis demonstrates significant reduction in the percentage (%) of TAMs (CD11b^+^ F4/80^+^), with a concomitant increase of **(I)** CD45^+^ TCRβ^+^ total T cells in the pancreatic tumors with LIF deletion, n=7-8 mice per group. **(J)** Flow cytometric analysis demonstrates a significant increase in the infiltration of CD45^+^ TCRβ^+^ CD8^+^ T cells, along with activated effector memory phenotype CD44^high^ CD62L^Low^, in tumors with cancer cell intrinsic deletion of LIF *in vivo* tumors as compared to LIF wild type, n=8 mice per group. **(K)** Schematic illustrating CREB-mediated transcriptional regulation of LIF expression, which polarizes macrophages toward an immunosuppressive phenotype via LIFR signaling, thereby facilitating crosstalk that attenuates T-cell infiltration and activation. Individual data points with mean ± SEM are shown and compared by two-tailed unpaired t test. *p<0.05; **p<0.01; ***p<0.001; ****p<0.0001; ^ns^ non-significant (p>0.05).

To achieve this, we polarized murine bone marrow derived naïve M0 macrophages (BMDMs) *in vitro* with conditioned media harvested from *KPC* PDAC cell lines (CREB^WT^ and CREB^KO^) (Supplementary Figure S4*A*). We then performed RNA-Seq on tumor-educated immunosuppressive macrophages (TEMs) to assess the role of tumor cell-specific CREB activation on their polarization state (Supplementary Figure S4*B*). DEG analysis comparing CREB^KO^ to CREB^WT^ TEMs revealed downregulation of several gene transcripts defining the M2-like, immune suppressive phenotype (*Tgfßi, Slc2A1, F13a1, Acly, Arg-1*) in CREB^KO^ TEMs, suggesting a role for tumor-specific CREB in altering macrophage polarization.

Next, we performed flow cytometric analysis on these BMDMs (Figure 5*B-D*). TEMs incubated with either recombinant (r) LIF or conditioned media harvested from CREB^WT^ PDAC cells showed a significant increase in the Arg-1^+^/iNOS^+^ ratio (gated within F4/80^+^ cells) as compared to M0 (Figure 5*B1*). Conversely, TEMs exposed to either CREB^KO^ alone or with LIFR blockade (EC359), with rLIF or CREB^WT^, not only led to a substantial reduction in this Arg1/iNOS ratio (Figure 5*B*) but also influenced reprogramming of these macrophages towards an antigen-presenting phenotype, as shown by upregulation of CD86 expression (Figure 5*C* and *D*). Taken together, these results suggest that LIFR blockade influences TAM reprogramming in CREB^WT^ conditions in a manner similar to CREB loss in *KPC* tumor cells.

Next, we investigated the direct impact of CREB-LIF conditioned BMDMs on T cell activation and function by isolating T cells from C57BL/6 murine spleen. T cells were cultured *ex-vivo* in the presence of conditioned media derived from polarized BMDMs [unstimulated (M0), M1 (LPS+ IFN-γ), M1+CREB^WT^ or CREB^KO^ or rLIF] in the presence or absence of EC359 (Supplementary Figures S4*C-E*). Suppression of T-cell IFN-γ secretion was observed following coculture of pan T cells with either rLIF or CREB^WT^ conditioned media (Supplementary Figures S4*D* and *E*). This suppression was rescued with CREB^KO^ alone or with LIFR blockade (EC359) with CREB^WT^, thereby strengthening the correlation of a CREB-LIF signaling axis in TAM-mediated immunosuppression and T cell activation *ex-vivo*.

To validate the role of LIF-LIFR signaling in shaping the TAM-mediated immunosuppressive TME, we employed the orthotopic *KPC* tumor model (Supplementary Figure S4*F).* Established tumors were treated with EC359 at a dosage of 15mg/kg for two weeks. Harvested tumors were then subjected to immunophenotyping analysis. Notably, we observed a significant reduction in Arg-1 and PD-L1 expression in intratumoral immunosuppressive macrophages from EC359 treated pancreatic tumors as compared to vehicle (Supplementary Figure S4*G)*. These changes in the innate immunosuppressive macrophage compartment correlated with a significant increase in the intratumoral expansion of activated effector (CD8^+^ CD69^+^ T cells) and effector memory (CD8^+^ CD44^High^ CD62L^Low^) T cells populations with EC359 treatment as compared to vehicle (Figure 5*E* and *F*).

To further substantiate that tumor cell-derived LIF acts via LIFR on macrophages to mediate these changes within the TME, we utilized lentiviral mediated CRISPR-*Cas9* to generate and expand clonal LIF knockout (LIF^KO^) *KPC* tumor cells (Supplementary Figure S4*H*). Cas9-expressing cells transduced with non-targeting small guide RNAs (sgRNAs) were used as controls (LIF^NTV^). LIF secretion as measured by ELISA was significantly attenuated in KO cells, confirming successful deletion in both clones (C-1 and C-6) as compared to control (Supplementary Figure S4*H*). LIF^KO^ (C-1) were used for subsequent experiments. Furthermore, compared to the control cells, LIF^KO^ *KPC* tumor cells demonstrated reduced expression of downstream effectors including stemness associated gene – *Cd44*, YAP1 signaling – *Cntf,* and JAK-STAT signaling components – *Stat3* (Supplementary Figure S4*I*), thereby providing further evidence for the disruption of tumor cell-intrinsic LIF dependent signaling in these KO cells.

Next, we generated tumors by implanting KPC-LIF^NTV^ or LIF^KO^ tumor cells into the pancreata of C57BL/6 mice (Figure 5*G*). Tumors were harvested for flow cytometric analysis 4 weeks after initial implantation. Compared with control, genomic deletion of LIF led to a significant decrease in the infiltration of innate Cd11b^+^F4/80^+^ macrophages (Figure 5*H*) with a concomitant increase in the frequency of total T cells (CD45^+^TCRβ^+^ T cells) and CD8^+^ T cells. Further, we identified enrichment of an effector memory phenotype (CD44^High^ CD62L^Low^) within these CD8^+^ T cells (Figure 5*I* and *J*). Collectively, these findings show a role for CREB regulated LIF in tumor cell-macrophage crosstalk within the TME, driving TAM-mediated T cell dysfunction (Figure 5*K*).

### Pharmacological targeting of CREB reduces pancreatic tumor burden and improves effector T cell infiltration in a murine model of PDAC

The functional redundancy observed between genomic deletion of *Creb* or *Lif* in attenuating the immunosuppressive tumor phenotype and augmenting effector T cell infiltration within the TME suggests that small molecule inhibitors capable of targeting CREB could be used to overcome dysfunctional T cell infiltration within the TME. We tested this hypothesis using a well-established small molecule inhibitor of CREB known as 666-15 (CREBi), which blocks CREB-dependent transcriptional activity and has been demonstrated to inhibit tumor growth in breast, pancreatic, and prostate cancer models (13, 24, 25). We first tested the therapeutic efficacy of 666-15 *in-vivo* in the orthotopic *KPC* mouse model (Figure 6*A*). At a dose of 20mg/kg, CREBi was found to inhibit tumor growth as compared to vehicle treatment (Figure 6*B*).

**Figure 6.**
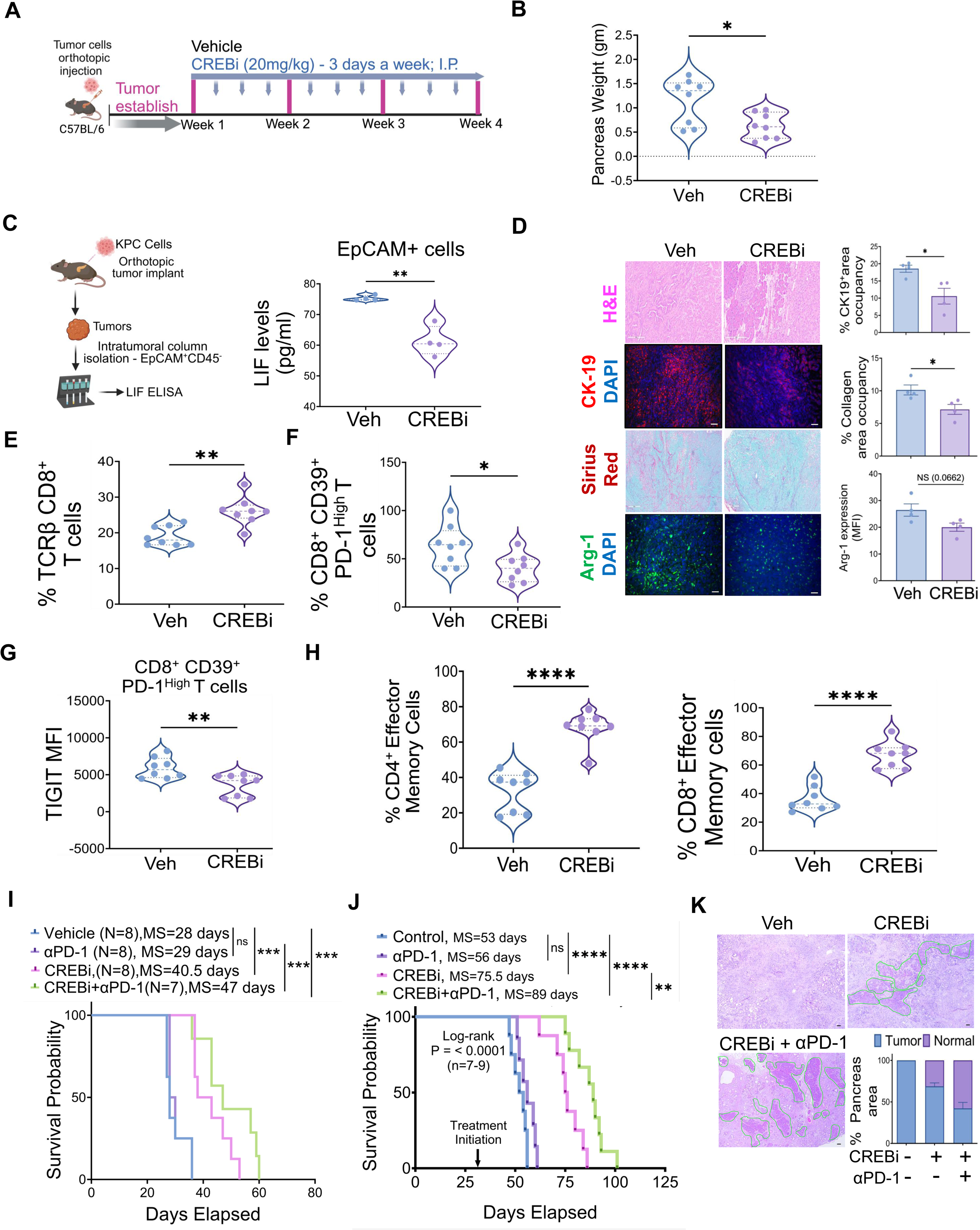
Therapeutic targeting of CREB with the small-molecule inhibitor (CREBi); 666-15 attenuates pancreatic tumor burden, reverses T-cell exclusion, and emerged as a potential therapeutic immuno-sensitizing strategy. **(A)** Schematic representation of the treatment regimen employed involving syngeneic orthotopic tumor cell implantation of KPC cells in C57BL/6 mice followed by i.p. administration of 666-15 (CREBi) at a dose of 20mg/kg for 3 weeks. Tumors were harvested on day 28 for downstream endpoint analysis. **(B)** Differences in pancreatic tumor weight between CREBi-treated and vehicle (veh) treated orthotopic tumors with n=8 mice in each arm. **(C)** Schematic representation (*Left*) along with violin plot depiction of LIF expression via ELISA (*Right*) in EpCAM^+^ sorted pooled single cell suspensions from orthotopic tumors treated with vehicle or CREBi. **(D)** H&E staining within the pancreas depicting poorly differentiated ductal adenocarcinoma in vehicle treated orthotopic tumors, whereas CREBi displayed higher presence of normal tissue architecture. Representative images along with corresponding quantification of CK19^+^, collagen (Sirius red), and Arg-1 positivity in the pancreatic tumor sections of vehicle or CREBi treated orthotopic KPC mice (n=4 mice in each group). **(E)** Flow cytometric analysis demonstrates a significant increase in the infiltration of CD45^+^ TCRβ^+^ CD8^+^ T cells, along with **(F-G)** reduction in the antigen experienced yet exhausted CD8^+^ CD39^+^ PD-1^high^ T cells expressing TIGIT with CREBi as compared to veh mice, n=8 mice per group. (**H)** Flow cytometric analysis demonstrates a significant increase in the infiltration of activated CD4^+^ as well as CD8^+^ T cells displaying activated effector memory phenotype (CD44^high^ CD62L^low^) within the pancreas of CREBi as compared to veh mice (n=8 mice per group). (**I)** Kaplan–Meier plot and log-rank test analysis showing improved overall survival of KPC orthotopic mice treated with CREBi alone or in combination with anti-PD-1 when compared with all other groups (n=7-8 mice per treatment arm). **(J)** Kaplan–Meier plot and log-rank test analysis showing improved overall survival of PKT orthotopic mice treated with CREBi alone or in combination with anti-PD-1 when compared with all other groups (n=7-9 mice per treatment arm). **(K)** Representative H&E-stained tumor sections along with quantification after 3 weeks of drug treatment depicting significant presence of normal pancreatic architecture (highlighted in green) in PKT mice treated with combination therapy of CREBi and αPD-1 as compared with veh or CREBi treated PKT (scale bar = 50 μm). Percentage of tumor area relative to the normal pancreas quantified within the whole pancreas, by capturing multiple fields of view. Individual data points with mean ± SEM are shown and compared by two-tailed unpaired t test for two group comparison. *p<0.05; **p<0.01; ***p<0.001; ****p<0.0001; ^ns^ non-significant (p>0.05).

Additionally, therapeutic targeting of CREB led to downregulation of LIF expression at both the transcript and protein levels (within the EpCAM^+^ cells) in these tumors (Figure 6*C* and Supplementary Figure S5*A*). Histological analysis of pancreata from CREBi treated mice revealed profound alterations in both the tumor cell and stromal compartments as shown in Figure 6*D*. H&E staining revealed a higher prevalence of normal pancreatic architecture correlating with attenuated disease burden along with a reduction in CK19⁺ ductal epithelial cells, decreased fibrosis (Sirius red), and a concomitant reduction in Arg-1 expression within the pancreata of CREBi-treated *KPC* mice compared to vehicle (Figure 6*D*).

Given the reduction in LIF-producing EpCAM^+^ cells after CREBi treatment and attenuation of stromal fibro-inflammation, we next examined whether CREBi impacts immunosuppressive myeloid-macrophage remodeling as well (Supplementary Figure S5*B*). Flow cytometric based analysis of CREBi treated pancreatic tumors revealed a significant reduction in Arg1 expression on TAMs (Gated within CD11b^+^ F4/80^+^ macrophages) as compared to vehicle. Notably, CREBi treatment also led to an upregulation of CD86 and PD-L1 expression on TAMs indicating a phenotypic shift towards a more immunostimulatory and potentially immune engaged phenotype (Supplementary Figure S5*B*). Furthermore, on examining the status of tumor infiltrating T cells in CREBi treated *KPC* orthotopic tumors, flow cytometric profiling revealed significant upregulation of CD8^+^ T cells with CREBi as compared to vehicle treatment (Figure 6*E*). Functionally, the proportion of antigen-experienced yet exhausted tumor-infiltrating PD1^+^CD39^+^CD8^+^ T cells (Figure 6*F*) harboring TIGIT expression was significantly reduced with CREBi treatment (Figure 6*G*). Importantly, the reversal of T cell dysfunction observed with CREBi was accompanied by a significant increase in the infiltration of CD4^+^ and CD8^+^ T cells with an effector memory phenotype (CD44^High^CD62^Low^) (Figure 6*H*). Taken together, these results suggest that CREB inhibition reinvigorates effector programs in tumor infiltrating T cells to augment anti-tumor immune response in PDAC, due, in part, to disrupting the tumor cell-TAM axis mediated by CREB-LIF.

### Therapeutic targeting of CREB in combination with PD-1 blockade reduces tumor burden in murine models of PDAC

Given the marked reduction in primary tumor burden, attenuation of fibroinflammatory milieu, and enhanced infiltration of activated effector CD8^+^ T cells following CREBi treatment, we next aimed to determine whether the addition of anti (α) PD-1 immunotherapy could enhance the therapeutic efficacy of CREBi in both aggressive orthotopic *KPC* and a highly aggressive preclinical GEMM of PDAC harboring oncogenic *Kras\** in a *Tgfbr2 –* null background. The *Ptf1a^cre/+^;LSL-Kras^G12D/+;^Tgfbr2^flox/flox^* (PKT) GEMM recapitulates key hallmarks of human disease, including pronounced stromal desmoplasia, immunosuppressive myeloid enriched TME, T cell exclusion, and, most importantly, lack of a therapeutic response to PD-1 blockade monotherapy starting at 4 weeks of age (13, 26–28). (Figure 6*I-J*). CREBi treatment significantly increased survival as compared to vehicle treatment [median survival (MS) 40.5 days vs 28 days in KPC and median MS 75.5 days vs 53 days in PKT]. Consistent with our previous studies (26) no difference in survival was observed with αPD-1 monotherapy when compared to vehicle (MS 29 days vs 28 days in KPC and MS 53 days vs 56 days in PKT model). The addition of CREBi to PD-1 blockade dramatically improved overall survival (MS 47 days vs 29 days in KPC and MS 53 days vs 89 days in PKT) as compared to αPD-1 monotherapy (MS 29 days in KPC and MS 56 days in PKT) or vehicle (MS 28 days in KPC and 53 days in PKT) treated *KPC* cohorts (Figure 6*I-J*). Moreover, analysis of the histological architecture in H&E-stained pancreatic tumor sections revealed a near 100% replacement of normal pancreatic parenchyma with tumor in vehicle-treated 4 weeks old PKT mice with substantial presence of late PanINs, cancer cells, and dense stromal architecture (Figure 6*K*). Notably, CREBi treatment alone or in combination with αPD-1 therapy resulted in a higher prevalence of normal acinar cells (highlighted in green), ADM, and early PanIN lesions, as compared to the vehicle cohort, providing additional validation of the profound anti-tumor response mediated by CREBi alone and in combination with PD-1 blockade in this highly aggressive PDAC GEMM (Figure 6*K*).

## Discussion

Oncogenic communication between cancer and immune cells within the TME plays a critical role in shaping the dynamics of PDAC and promoting therapeutic resistance. The present study illustrates a previously unrecognized role of cancer cell-intrinsic activation of CREB in coordinating cancer cell – macrophage communication in shaping the immunosuppressive TME which mediates T cell exclusion and poor response to immunotherapy in PDAC. Mechanistically CREB deletion in tumor cells reprograms immunosuppressive macrophages towards an immune-stimulatory phenotype, enhances effector T cell infiltration, and attenuates disease progression. At the molecular level, we uncovered CREB-dependent transcriptional control of LIF, which mediates paracrine crosstalk with LIFR-expressing macrophages, driving their polarization towards an immunosuppressive state putatively by activating the JAK-STAT3 signaling axis. Therapeutic targeting of CREB using a small molecule inhibitor recapitulates the genetic phenotype affiliated with CREB loss and, importantly, synergizes with PD-1 checkpoint inhibition to produce anti-tumor responses in otherwise refractory PDAC models. Overall, these findings position CREB as a pivotal node linking cancer cell autonomous oncogenic signaling to macrophage driven immune suppression, offering rationale for therapeutic targeting of the CREB-LIF axis to restore anti-tumor immunity in PDAC. Although, these results highlight CREB as a therapeutically actionable hub converging on oncogenic and immunomodulatory programs, targeting transcription factors has historically been challenging due to multiple off target effects and toxicity profiles (29). Given its central role in regulating a myriad of cellular stress responses, metabolic adaptations and neuronal signaling (30), these concerns highlight the need for more innovative and precision therapeutic strategies, that selectively modulate oncogenic CREB activity in tumor cells while sparing homeostatic CREB functions in non-malignant tissues intact.

While most prior studies have focused on the role of CREB in regulating cancer-cell autonomous mechanisms of oncogenic growth (25), our findings reveal a novel paradigm in which CREB directly engages in distinct paracrine interactions with TAMs within the TME of PDAC. Physiologically, macrophages are central mediators of tissue homeostasis, but in tumors their functional plasticity is co-opted to promote cancer progression (31). TAM activation states exist along a continuum of phenotypes which are often dictated by development of origin, tissue context and cues from the local microenvironment (32). Although our understanding of TAM biology and heterogeneity has increased considerably over the years, the distinct paracrine mechanisms driven by cancer cell intrinsic CREB signaling that influence macrophage biology have not been reported previously. Our present data in CREB deleted-PDAC tumors identifies a significant reduction in immunosuppressive polarized pro-tumor TAMs expressing high Arg1, MRC1, STAT3, accompanied by increased MHC class II antigen presentation and enrichment of the T cell–recruiting chemokines CXCL9 and CXCL10. High Arg1 expression has been well established as a functional marker of immunosuppressive myeloid cells, that suppressed the T cell based anti-tumor response by depleting the amino acid arginine (33). Consistent with functional reprogramming of TAMs *in vivo* with CREB deletion, our results also demonstrate heightened infiltration and activation of T cells providing additional evidence for a polarized shift towards an immunostimulatory TME with CREB loss.

In parallel, to dissect CREB-mediated mechanisms governing TAM polarization and function, our unbiased RNA transcriptomic analysis implicated LIF, a pleiotropic immunomodulatory cytokine of the IL-6 superfamily previously implicated in TAM-mediated immunosuppression and T cell exclusion (22, 34), as a putative downstream target of CREB that may mediate these processes, thereby uncovering a novel regulatory axis in LIF biology. Prior scientific evidence highlights a prominent role of LIF overexpression in solid tumors including PDAC as a marker for poor prognosis and resistance to conventional chemo or immunotherapy (23, 34, 35). Consistent with its previously reported role in driving TAM-mediated immunosuppression and restricting effector CD8^+^ T-cell infiltration into the TME, our study also firmly establishes that pharmacological blockade of LIFR using a small molecule inhibitor (EC359) or genetic ablation of LIF effectively reverses this immunosuppressive TAM phenotype and overcomes T cell exclusion within the PDAC TME.

While our previous work demonstrated that CREBi alone can reinvigorate the TME of pancreatic tumors in an immunologically inert, highly stromal desmoplastic and T cell excluded PKT GEMM of PDAC (13), the present study advances this knowledge by elucidating tumor stromal and immune crosstalk mechanisms that had remained unexplored. We demonstrate that CREB not only impacts cancer cell dynamics but also the stromal and immune microenvironment as well. While systemic inhibition of CREB may influence a plethora of secreted disease factors communicating across different cell types, a recent study highlights a role of LIF as one of the major mechanisms of paracrine communication between cancer associated fibroblasts (CAFs) and cancer cells modulating therapeutic response in PDAC (36). To this end, our data suggest that CREB inhibition mitigates stromal inflammation by attenuating LIF expression, putatively through both autocrine and paracrine mechanisms.

PDAC tumors are notoriously adept at employing diverse strategies to establish a protumorigenic and immunosuppressive TME. Several of such adaptive mechanisms by which cancer cells evade immune surveillance involve stromal desmoplasia, including the presence of TAMs highly expressing Arg-1 along with secretion of cytokines including IL-6 and TGF-β. Together, these factors contribute to either the exclusion of T cells or the suppression of effector T cell function, both of which are recognized as critical drivers of therapeutic resistance. In addition to mitigating stromal inflammation, our present study demonstrates CREBi promoted the reprogramming of TAMs towards a more immunostimulatory phenotype, characterized by elevated PD-L1 and CD86 expression. Consistent with this, a recent study by Wang et al. demonstrated that PD-L1⁺ TAMs are more mature and activated, capable of sustaining CD8⁺ T cell proliferation and cytotoxic activity, and that their higher density correlates with improved prognosis in breast cancer (37).

Our data further highlights that overcoming these barriers of therapeutic resistance can alleviate T cell exclusion and reinvigorate the TME with heightened effector T cell infiltration. While the promise of immunotherapy has profoundly altered the clinical outcomes of many cancer types, this paradigm shift has not extended to PDAC patients (9). Concordant results in preclinical PDAC models implicate the stromal-immune microenvironment involving dysfunctional T cells and presence of TAMs as the critical molecular determinants of this phenotype. Intriguingly, our flow cytometric analysis in CREBi treated mouse tumors demonstrated a significant reduction in the antigen experienced yet exhausted CD8^+^CD39^+^ T cells harboring TIGIT expression. CD39 is an ectonucleotidase expressed on exhausted or dysfunctional T cells within the tumor (38, 39). With its ability to deplete extracellular ATP, high CD39 enzymatic activity results in generation of AMP resulting in inadequate T cell activation and dysfunctional CD8^+^ T cells expressing inhibitory checkpoint receptors (40). By reversing this exhaustion phenotype, CREBi not only restores effector CD8⁺ T cell functionality but also enhances the quality of adaptive immune cells within the pancreatic tumor milieu. These findings provided us with a compelling rationale to explore combinatorial strategies of CREBi with anti PD-1 therapy in preclinical murine models of PDAC.

In summary, our study provides critical insights into the functional and preclinical significance of cancer cell-autonomous CREB expression in shaping the immune microenvironment through LIF signaling. Findings from CREB-deleted PDAC models reveal that attenuation of CREB-dependent transcriptional programs reinvigorates an immunostimulatory TME, at least in part due to blocking paracrine LIF engagement with LIFR on TAMs. This improves T cell infiltration/function and sensitizes otherwise immunorefractory PDAC to anti PD-1 therapy. Prospective validation and clinical translation of CREB/LIF-LIFR targeted therapies are warranted in future studies.

## Materials and Methods

### Cell Lines

Human pancreatic cancer cell lines (Panc02.13, Panc08.13, MiaPaCa, Panc1, HPAC, BxPC-3), non-malignant epithelial (HPNE) cells along with murine RAW 264.7 cells (Macrophage) line were obtained from ATCC (https://www.atcc.org/). Immortalized murine PDAC cells (K8484, and KPC-6694) and were obtained, established and maintained as described previously (41, 42). Cells with low passage numbers (<10) were used in the study and tested for detection of Mycoplasma contamination using the MycoStrip kit (#rep-mys-50, InvivoGen) (March 2023).

### Histological Analysis and Quantification

Pancreas tissue sections were fixed in 10% neutral-buffered formalin, embedded in paraffin, and sectioned. Subsequent histological staining techniques, including Hematoxylin and eosin (H&E), Sirius Red, and Alcian Blue, were performed following established protocols as described (15, 17, 26, 43). Following deparaffinization, antigen retrieval and quenching with endogenous peroxide, the sections were then incubated overnight at 4℃ in a humidified chamber with primary antibodies listed in Supplementary Table S1. Following washing, the sections were developed using the Vector^®^ ABC Kits (HRP-based kit SK-Vector Laboratories) according to the manufacturer’s instructions, with diaminobenzidine (DAB) serving as the chromogen. For dual color IHC, tissue sections were stained using violet (purple) reaction product as a second color for multiple antigen labelling using Vector^®^ VIP Substrate Kit, Peroxidase (HRP). Mayer’s hematoxylin was used for counterstaining, and images were acquired using a DM750 Leica microscope (Leica Microsystems). Image J based quantification was used to depict percentage area occupancy, images were first converted into grayscale and set at a threshold to calculate collagen positive area accumulation for Sirius red. For DAB-stained mouse IHC tissue sections, ImageJ color deconvolution plugin to isolate DAB from hematoxylin, and the percentage of + area was calculated across multiple pancreatic fields of view per mouse sample, using consistent thresholds to minimize background noise.

### Human PDAC tissue Microarray

Commercially purchased tissue microarray (TMA) slides (Cat. No. PA2081c, PA2082as TissueArray.Com LLC), which contained tissue cores representing a range of pancreatic disease states. PDAC samples were subjected to staining for phosphorylated CREB (pCREB) using single-color IHC or dual color pCREB/CD68 staining. Tissue staining was evaluated using a staining index, calculated as the sum of intensity scores, following the guidelines provided by an expert pathologist as described in detail previously (13). The staining index was calculated as the sum of intensity score (0, no staining; 1+, weak; 2+, moderate; 3+, strong) with percent distribution (0, no staining; 1+, staining of <33% of cells; 2+, between 33% and 66% of cells; and 3+, staining of >66% of cells). Staining indices were classified as follows: 2+ to 3+ or higher, = strong staining; 0 to 1+, = weak staining.

### PDAC Datasets Expression Analysis

PDAC (TCGA) and normal pancreatic tissue (GTEx) gene expression data were retrieved from the Gene Expression Profiling Interactive Analysis (GEPIA2). Expression analysis was conducted to compare CREB mRNA expression across classical (N=86) and basal (N=65) PDAC subtypes to healthy (N=171) pancreatic tissues. Statistical significance was assessed using ANOVA test with p-values less than 0.05 considered significant. *CREB* mRNA expression was plotted against progression free survival TCGA PDAC (PAAD) patient cohort on cBioPortal platform (44–46). Similar steps outlined were followed to depict LIF expressions in PAAD vs normal pancreas datasets using GEPIA2 as well. For immune cells deconvolution analysis, relative abundances of 22 immune cell types were estimated for each of TCGA-PAAD samples using CIBERSORT based on LM22 signatures (containing 547 genes) (47).

### Human Tumor Atlas Network (HTAN) Datasets Analysis

Single-cell RNA sequencing (scRNA-seq) data from 7 PDAC patients were obtained from the Human Tumor Atlas Network (HTAN) repository (48), deposited by the Washington University Human Tumor Atlas Research Center. A total of 25 samples (from 7 patients) were analyzed using Seurat (v5.1.0) (49). Downstream analysis was conducted in R (v4.3.3) using the Seurat package (v5.1.0). Low-quality cells were excluded based on thresholds for mitochondrial gene content (>10%), unique gene count (<500 or >7,000), and total UMI count (<60000). Data from all samples were merged into a single Seurat object and normalized using SC Transform (v2), with default settings for regression and variable feature selection. Dimensionality reduction was performed via PCA followed by UMAP embedding. Clustering was performed using the Louvain algorithm with resolution tuned empirically with resolution tuned empirically using the Cluster R package to identify a stable clustering structure. Batch correction and dataset integration were executed using Harmony (v1.2.3). Following integration (Total cells-153033), cluster annotation was performed using canonical marker genes identified with FindAllMarkers (minimum percentage expression = 0.1; positive log-fold change; adjusted p-value < 0.05). All statistical analyses were performed in R (v4.3.3), and visualization was conducted using ggplot2 (v3.5.1), and patchwork (v1.3.0) packages.

### Defining High and Low CREB1 Expression using GSEA analysis in scRNA seq HTAN datasets

Gene expression data was utilized from tumor cells cluster from HTAN data. A curated gene set associated with CREB1 signaling from the MSigDB (50) human gene set was used, including the following genes-PIK3R1, PIK3CA, AKT1, MAPK3 and CREB1. Duplicate genes were removed to ensure data integrity. ssGSEA was performed using the *gsva* function from the GSVA R package (51), gene set enrichment scores were calculated that reflected CREB1 pathway activation in each cell type. The following parameters were used: method = “ssGSEA”, min.sz = 1, max.sz = 1000. Median ssGSEA score was used as a cutoff to classify tumor cells by CREB1 activity, into “High” and “Low” CREB1 activity groups. These classifications were used for further downstream analysis.

### Cell Chat DB Analysis

Following ssGSEA classification of tumor cells, CellChat (52) analysis was performed to investigate the intercellular communication networks in the tumor microenvironment. Cell-cell communication network was constructed using relevant ligand-receptor pairs within the tumor cell cluster. Communication strengths between different tumor cells were calculated, and differential patterns were compared between the “High” and “Low” CREB1 activity groups. Communication networks were visualized using CellChat network plots.

### IF and RNA fluorescence in situ hybridization (FISH) imaging

IF based imaging was conducted on paraffin embedded tumor tissue sections. These sections were subjected to deparaffinization, permeabilization followed by heat induced antigen retrieval using citrate buffer (0.01 M, pH 6.0) and staining with primary antibodies. Fluorescent-labeled secondary antibodies were appropriately diluted in Phosphate Buffered Saline (PBS, 1X, Gibco) at 1:1000 ratio and applied to slides for 30 minutes at room temperature. Slides were sequentially washed three times with PBS. of Hoechst 33342 dye at 20 μg/ml was employed for nuclear staining. Multiple fields of view were acquired at 10x or 20x magnification using the DMi8 fluorescent microscope (Leica Microsystems) or Dragonfly high-speed confocal (Andor, Oxford Instruments). For IF based staining, primary antibody was detected using species-specific Alexa Fluor 594 and/or Alexa Fluor 488 (Thermo Fisher) secondary antibodies. Image J based quantification of IF images were performed using digital image conversion to 16-bit grayscale followed by threshold adjustment to distinguish true signals from background autofluorescence. Analyzing particle’s function was used for automated cell counts in Image J to depict Ki67^+^ cells.

Customized HCR^TM^ murine RNA-FISH probe sets against *Creb1* mRNA with appropriate amplifiers were purchased directly from the Molecular Instruments (Los Angeles, CA). Initial tissue processing for Co IF + RNA-FISH was carried out in the same fashion as IF mRNA detection and amplification stages were performed using Molecular Instruments HCR^TM^ IF+ HCR^TM^ RNA-FISH as per manufacturer protocol.

### Animal Studies

Mice of both sexes weighing 20 to 25 g were used; they were housed in pathogen-free conditions under a 12-hour light-dark diurnal cycle with a controlled temperature of 21^0^ C to 23^0^C and maintained on a standard rodent chow diet (Harlan Laboratories) before the experimental induction including orthotopic mice surgeries. The mice were euthanized upon the manifestation of signs of compromised health, including weight loss, accelerated respiration, hunched posture, piloerection, and reduced activity. All animal experiments were approved and performed in compliance with the regulations and ethical guidelines for experimental and animal studies of the Institutional Animal Care and Use Committee and the University of Miami guidelines (Miami, FL; Protocol Nos.18-081, and 21-093).

### Creb Deletion in a Genetically Engineered Kras^LSL-G12D/+^; Trp53^LSL-R172H/+^ Pdx1^Cre/+^; (KPC) Mice Model

*Pdx1^Cre^*^/+^ knock-in allele mice were obtained from Jackson Laboratory (Bar Harbor, ME; strain number: 014647). *Creb^fl/fl^*mice were obtained from Professor Eric Nestler (Cold Spring Harbor Laboratory, Cold Spring Harbor, NY 11724) and were crossed with *Pdx1^Cre^*^/+^ to obtain *Pdx1^Cre^*^/+^*Creb^fl/fl^* mice. To generate *Pdx1^Cre^*^/+^ *Kras^LSL-G12D/+^*; *Trp53^LSL-R172H/+^; Creb^fl/fl^*(*KPCC^-/-^*), each of the transgenic mice, i.e. LSL *Kras^G12D/+^*; LSL *Trp53^R172H/+^*; and *Pdx1^Cre^*^/+^ were individually crossed with *Creb^fl/fl^* to generate each of the transgenic mice in *Creb^fl/fl^* background (LSL-*Kras^G12D/+^/ Creb^fl/fl^*, LSL-*Trp53^R172H/+^*/*Creb*^fl/fl^, and *Pdx1-Cre^/+^/Creb*^fl/fl^). Thereafter, *LSL-Kras^G12D/+^*/*Creb*^fl/fl^ mice were crossed with LSL-*Trp53^R172H/+^*/*Creb*^fl/fl^ to get *KPC^-/-^* (LSL-*Kras^G12D/+^;* LSL-*Trp53^R172H/+^*; *Creb^fl/fl^*) mice. Finally, *KPC^-/-^* mice were crossed with *Pdx1-Cre/Creb^fl/fl^* mice to get LSL-*Kras^G12D/+^*; *LSL-Trp53^R172H/+^*; *Pdx1-Cre; Creb^fl/fl^* (*KPCC^-/-^)* mice. All mice were housed under standardized conditions at the University of Miami animal facility. All *in-vivo* mice related experimental procedures were reviewed and approved by the University of Miami Institutional Animal Care and Use Committee (IACUC, protocol #18-081, 21-093) in accordance with institutional guidelines.

### Mouse Genotyping

Genotyping was conducted using the automated genotyping service provider, Transnetyx. Supplementary Table S2 presents the probe sequences used for genotyping analysis.

### H&E-based Assessment of Pancreatic Lesions

Pancreatic tissue sections from age-matched *KC*, *KPC* and *KPCC*^-/-^ GEMM in each group were stained with H&E and subsequently examined for ADMs, pancreatic lesions, and carcinoma in situ. The proportion of acinar area and the number of ducts harboring PanINs (any grade) were quantified by surveying multiple non overlapping fields of view. PanINs were graded according to the established criteria (53, 54) and as also described in detail in our recent publication (17).

Quantitative histological analysis in PKT GEMM pancreatic tumor tissue sections was conducted by evaluating the fraction of tumor area for each treatment cohort and as described in detail previously (26, 41). To analyze the percentage of the tumor area in relation to the entire pancreas, we captured multiple fields of view (3–4) at lower magnification. We quantitatively compared total area of normal pancreatic architecture (without Pancreatic Intraepithelial Neoplasia (PanINs) and tumor ductal epithelial cells) with areas with tumor cells in each tissue section.

### Flow Cytometry

Mice pancreatic tumors were enzymatically digested to obtain single cell suspensions as described in detail previously (15, 26). Samples were washed with autoMACS rinsing buffer + BSA (Miltenyi Biotec, #130-091-222, #130-091-376), incubated with FcR-blocking antibody (Miltenyi Biotec, #130-092-575), and subsequently stained with fluorescently conjugated antibodies (Supplementary Table S3). LIVE/DEAD^TM^ Blue Fixable (Invitrogen, #L23105) is used to discriminate live/dead cells as per manufacturer’s protocol. Cells were fixed and permeabilized using FoxP3 staining buffer kit (Thermo Fisher Scientific, #00-5523-00). Flow cytometry data acquisition was performed on Cytek Aurora and analyzed using FlowJo v.10 software. Gating strategies depicting myeloid macrophages and T cell subsets are depicted in Supplementary Figure S6.

### Orthotopic Tumor Cell Implantation and In-vivo Treatment in Genetically Engineered Mouse Models of PDAC

Single cell suspension of murine PDAC cells were prepared in DMEM, washed with sterile PBS and resuspended with matrigel and were injected orthotopically directly into the pancreatic tail of C57BL/6 mice with 10 ul final volume per mouse as described in detail previously (26). Prior to implantation the cells were confirmed to be free of any pathogenic microbial contamination using standard sterility assays. Both male and female mice were used for orthotopic tumor implantation in the present study. Once the tumors were established following initial implantation, mice were randomized to receive intraperitoneal (i.p.) injection of CREB inhibitor (CREBi, 666-15, 20mg/kg) [MedChemExpress, #HY-101120] administered 3x weekly, αPD-1 antibody (BioXCell, Clone #BE0273, 200 μg/mouse, intraperitoneal injection twice weekly), or a combination of CREBi and αPD-1 until moribund for survival studies. EC359 [MedChem Express, HY-120142] is a potent inhibitor of leukemia inhibitory factor receptor (LIFR) were administered subcutaneously at a dose of 15 mg/kg, thrice weekly for a total of 2 weeks. Animals showing clinical signs of significant disease burden (impaired mobility, ruffled hair, hunched posture, loss of coordination) were euthanized. For endpoint-based analysis in *Ptf1a^cre/+^;LSL-Kras^G12D/+^;Tgfbr2^flox/flox^* mice (PKT), both male and female mice at 4 weeks of age were treated for 3 weeks with CREBi (10mg/kg, i.p. administered 5 days on/2 days off), αPD-1 antibody (200 μg/mouse, i.p. twice weekly) or the combination. Pancreatic tumors were harvested at the end of treatment duration for downstream analysis.

### Nanostring Digital Spatial Profiler in KPC GEMM of PDAC

Pancreatic tumor sections from age matched *KPC* and *KPCC^-/-^ GEMM* (2 biological replicates per group) were stained with cell morphology markers (Green, pan-cytokeratin), TAMs (Red, F4/80) and a nuclear DNA (Blue, SYTO13) stain. Stained slides were subsequently scanned using GeoMx^TM^ Digital Spatial Profiler platform (DSP). Following Nanostring best practices and standardized workflows, regions of interest (ROIs) were selected and segmented, ensuring a minimum of 20 cells per area of illumination (AOI), to distinguish macrophage and tumor cell compartments. Next, AOIs were subjected to UV illumination to photocleave the oligonucleotide tags from the bound protein probes. The released tags were collected into designated wells, hybridized and digitally counted on the DSP. Raw counts were processed with standard QC including negative control (IgG) background assessment and normalized using Q3 and/or housekeeping proteins. Features below the limit of quantitation were excluded before downstream differential expression analysis.

### Sample preparation for scRNA sequencing in orthotopic PDAC tumors

Pancreas tumor tissue harvested from mice bearing orthotopically implanted CREB^WT^ and CREB^KO^ (n=3 mice per group, pooled). Tumors were mechanically dissociated to generate single cell suspensions as described previously. Cells were then resuspended in 10 mL of buffer (autoMACS Rinsing Solution in 5% MACS BSA) and incubated for 10 minutes at 4 °C with 4′,6-Diamidino-2-phenylindole dihydrochloride (Roche Diagnostics) 1:1000 and DR ™ (Biolegend) 1:500 to stain cells for viability. Live cells sorted by BD FACS Aria Fusion were then resuspended in PBS (BioWhittaker, Inc) with 3% BSA (5% MACS BSA) for single-cell RNA sequencing. Single cell RNA sequencing the 10x Genomics Chromium Single Cell 3’ Reagent v3.1(# PN-1000268) was used with standard conditions and volumes to process cell suspensions for 3’ transcriptional profiling. Target cell recovery, cDNA library preparation, and subsequent quantitative and qualitative assessments were performed following the procedures outlined in our earlier publications (26). Raw intensity files were demultiplexed into FASTQ format using Illumina Base Space software followed by aligning to the transcriptome using the 10x Genomics Cell ranger (v4.0.0) package.

### Cluster Identification and Annotation of ScRNA Sequencing Dataset

Single cell gene expression matrices were processed using Seurat (v4.3.0). Cells fewer than 200 detected transcripts and those exhibiting mitochondrial content above 7.5% were excluded. Gene expression counts were normalized with the *NormalizeData* function applying a scaling factor of 10,000 and the *LogNormalize* method. Highly variable genes were identified using the *Find Variable Features* function. Data were subsequently scaled and centered through linear regression on the counts and the cell cycle score difference. Principal Component Analysis (PCA) was performed using *RunPCA* function using the previously defined variable genes to identify necessary dimensions for >90% variance within the data. Violin plots were then used to filter the data according to user-defined criteria. Cell clusters were obtained via the *Find Neighbors* and *FindClusters* functions, using a resolution of 0.7 and non-linear dimensional reduction was then performed using Uniform Manifold Approximation and Projection (UMAP) clustering. Differentially expressed features across clusters were identified with FindAllMarkers, and clusters were annotated according to marker gene expression. Clusters displaying overlapping expression of established lineage specific markers were merged for downstream analysis.

### Differential Gene Expression Analysis of Single-cell RNA Sequencing Dataset

To assess differential gene expression analysis, each cluster of interest was first subsetted, re-normalized, scaled and subjected to PCA. Differentially expressed genes (DEGs) between identity classes were identified using “*FindMarkers*” function from Seurat v4.0 as described in detail previously (17).

### Western Blot Analysis

The freshly harvested mouse or human cell lines were promptly flash-frozen and preserved at - 80°C for long-term storage. Frozen cells pelleted were thawed and homogenized in RIPA buffer (0.1% SDS, 50 mM Tris·HCl, 150 mM NaCl, 1% NP-40, and 0.5% Na deoxycholate) with protease inhibitor cocktail (# P2714, Sigma, St. Louis, MO) and PhosSTOP phosphatase inhibitor ((#04906845001, Roche, Indianapolis, IN, USA). Samples were sonicated for 2 minutes on ice, centrifuged at 10,000 g for 15 minutes at 4^0^C to collect supernatant. Supernatants were then collected, quantified with BCA assay (Thermo Fisher Scientific, 23227). Lysates containing equal protein were separated using 4-20 % SDS PAGE Mini-Protean TGX Stain-Free Gel (Bio-Rad, #4568096) and transferred on Trans-Blot® Turbo™ Midi PVDF Transfer Packs (Bio-Rad, #1704156EDU) using Trans-Blot Turbo Transfer System (Bio-Rad). For immunodetection, membranes were incubated with primary antibodies listed in Supplementary Table S1 at 4°C overnight. Membranes were then washed and incubated with corresponding specific secondary antibodies conjugated with horseradish peroxidase (Jackson ImmunoResearch Laboratory). Immunoreactive bands were developed using Pierce ECL Western Blotting Substrate (Thermo Fisher Scientific, #32106) or SuperSignal™ West Pico PLUS Chemiluminescent Substrate (Thermo Fisher Scientific, #34580). Uncropped images of the blots are shown in Supplementary Figure S7.

### RNA Isolation and qPCR Analysis

Total RNA was extracted from flash-frozen pancreatic tissue or cultured cells using the RNeasy Kit (#74134, Qiagen) following the manufacturer’s instructions. cDNA was synthesized by reverse transcription of the isolated RNA and subsequently used for quantitative PCR (qPCR) analysis using gene-specific predesigned primers (RT^2^ qPCR Primer Assay, Qiagen listed in Supplementary Table S4) and iQ^TM^ SYBR^®^ Green Supermix (#1708880, BioRad). Gene expression levels were normalized to the housekeeping mouse 18s rRNA gene using the comparative CT (^ΔΔ^CT) method and reported as fold change relative to control.

### Statistical Analysis

Descriptive statistics were calculated using Prism software (GraphPad Software Inc). Results are shown as values of means ± SEM unless otherwise indicated. Prior to applying any parametric tests, data distributions were evaluated for normality using Shapiro-Wilk test. For experiments involving more than two groups, one way ANOVA was performed followed by Tukey’s or Games-Howell post hoc tests as appropriate. A two-tailed Student’s t test was used for two group comparisons. Statistical significance was defined using a cutoff of 0.05, except when indicated in the figure legend otherwise. Pairwise comparison of survival curves between individual treatments was performed using the log rank (Mantel-Cox) test. Differences in the distribution of IHC staining categories between groups were evaluated using Fisher’s exact test.

### Chromatin Immunoprecipitation (ChIP) sequencing

Formaldehyde fixed KPC tumor cells were prepared as per the Active motif’s (Carlsbad, CA) protocol. Briefly, 5 million cells were fixed in formaldehyde for 15 minutes followed by incubation in glycine solution for 5 minutes to stop the fixation. Then, cells were centrifuged at 800 x g for 10 minutes to remove the supernatant followed washing with chilled PBS-glycine and 100 µl 1mM PMSF. The final pellet was shipped in dry ice to Active Motif for the subsequent steps (chromatin preparation, chromatin immunoprecipitation using total CREB1 antibody, library generation, sequencing and data analysis) of ChIP-sequencing. The ChIP-seq integrated genome viewer seq tracks were generated using UCSC genome browser interface using bigwig files provided by Active motif.

### CRISPR/Cas9 Gene Editing in KPC PDAC tumor cells

Genetic ablation of mouse CREB in KPC PDAC cells were achieved using transduction of commercially purchased viral particles containing TransEDIT mouse CRISPR gRNA target gene set for *Creb1* (CCMV1101-12912, Transomic Technologies, Huntsville, AL) along with non-targeting control viral particles. In brief, 7 x 10^4^ target cells were transduced using 5-8 μg/ml Polybrene with viral particles in line with the technical manual of transEDIT Lentiviral gRNA plus Cas9 (pCLIP-All) Target Gene Sets (Transomic Technologies, Huntsville, AL) in a 12 well plate. Cells were incubated for 12-24 hours and then replaced by fresh media. After 24-48 hours transduced cells were selected using fluorescent protein expression. Later, using fluorescence based activated cell sorting analysis (FACS), single cell clonal population were generated and expanded into 96 well plates in 100 μl of complete media. Individual colonies were expanded and validated (KO) of respective protein by western blot analysis for total CREB protein (48H2, CST#9197S, Cell Signaling Technology, Danvers, MA).

Oligos of sgRNAs targeting mouse *Lif* were designed by CRISPick Broad Institute (Sequence of oligos listed in Supplementary Table S5). Genetic Perturbation platform, the backbone vector LentiCRISPRv2-mCherry (a gift from Agata Smogorzewska [Addgene plasmid # 99154; http://n2t.net/addgene:99154;RRID:Addgene_99154] that confers sgRNA cloning site + hSPCAS9-P2A-mCherry were used to clone and ligate these guides. These constructs were packaged as lentiviruses using third generation lenti virus packaging systems with HEK293T cells standard protocols. The lentiviral particles generated in HEK293T packaging cells were used to transduce mouse KPC cells with polybrene, following recovery, these cells were subjected to FACS to isolate mCherry^high^ population. Sorted cells were maintained in complete DMEM containing 10% FBS and subsequently plated at ultra-low density in 150-mm culture dishes. Individual colonies were manually picked under a microscope and clonally expanded. Successful knockout was then validated by assessing LIF protein levels in the conditioned media using ELISA.

### Isolation, Differentiation and Polarization of Bone Marrow Derived Macrophages (BMDMs)

Bone marrow from long bones (tibia, femur) was isolated from C57BL/6 mice and maintained in RPMI with 10% FBS, penicillin/streptomycin (Gibco) for 5 days with M-CSF (Biotechne) at a concentration of 30ng/ml. Media was changed on day 3, naïve macrophages were polarized on day 5. BMDMs were polarized with either 20 ng/ml murine LPS (Biotechne) or IFN-γ (Biotechne) and murine IL-4 (20 ng/ml) or conditioned media derived from KPC CREB^WT^ or CREB^KO^ tumor cells. To assess the polarization pattern of bone marrow derived macrophages, these cells were maintained in these respective experimental conditions for at least 48 hours. Under specific experimental conditions, bone marrow derived macrophages (BMDMs) were treated with murine recombinant LIF (Biotechne, 1 µg/mL), with or without pretreatment using an LIF receptor (LIFR) inhibitor (0.2 µM EC359, HY-120142, MedChem Express) at 37°C for 30 minutes. Cells were then harvested and subjected to downstream assays.

### Bulk RNA Sequencing

Total RNA isolated from each (3–4) technical replicates were submitted to the Onco-Genomics Shared Resource (University of Miami Miller School of Medicine, Miami, Florida) for library construction. RNA-seq libraries were prepared using the TruSeq® Stranded Total RNA Library Prep Gold (Illumina, 20020599). In brief, extracted RNAs were quantified and qualitatively assessed on the ThermoFisher QuBit Fluorometer’s High Sensitivity RNA (Cat #Q32855) and Agilent Fragment Analyzer High Sensitivity RNA kits (Cat #DNF-472-0500), and up to 500 ng of RNAs were used as input. Total RNA was first depleted of ribosomal RNA using RiboZero beads, followed by fragmentation, then first and second strand cDNA conversion, and finally library synthesis with 3’ adenylation, adapter ligation and PCR amplification. Finished libraries quantified and qualitatively assessed on the ThermoFisher QuBit Fluorometer’s High Sensitivity DNA (Cat #Q32855) and Agilent Fragment Analyzer High Sensitivity NGS kits (Cat # DNF-474-0500) prior to normalization and pooling in preparation for sequencing. Sequencing was performed at the OncoGenomics Shared Resources (OGSR) of Sylvester Comprehensive Cancer Center. In brief, pooled libraries were sequenced using single-end or paired-end 101 base pair chemistry on an Illumina NextSeq500 or NovaSeq6000 instrument. Targeted average sequencing depth for RNA-seq was between 20 and 40 million clusters per library.

### Enzyme-linked immunosorbent assay (ELISA)

Conditioned medium was collected from cells, cleared by centrifugation, and analyzed using a commercially available sandwich ELISA kit for mouse LIF (# MLF00; R&D systems), mouse IFN-γ (# MIF00-1; R&D systems) according to the manufacturer’s instructions. Bulk tumor and cell lysates were quantified and 300-400 ug of protein was loaded per well. All ELISAs were performed with 3-4 technical replicates. For quantification of LIF in orthotopic KPC PDAC tumors, CD326^+^ (EpCAM) cells were isolated by using CD326^+^ microbeads (Miltenyi 130-105-958), a MIDIMACS separator and LS columns (#130-042-401). Levels of mouse LIF were estimated using pooled EpCAM^+^ cell lysates (n=5 mice per group).

### T cell coculture with TAM Functional Assay

Splenic T cells were isolated from C57BL/6 mice using a Pan T Cell Isolation Kit (Miltenyi Biotec, #130-095-030) according to the manufacturer’s instructions. The isolated T cells were cultured in complete media and were activated using Rapid T cell activation kit (#86772, Cell Signaling Technology). In parallel, pancreatic single cell suspensions prepared from CREB^WT^ (n=4) or CREB^KO^ (n=4) tumors were used to isolate F4/80^+^ TAMs using anti-F4/80 microbeads ultra mouse kit (#130-042-401, Miltenyi) as per the manufacturer’s protocol. At the final step, F4/80^+^ cell pellet from biological replicates of respective mice cohort were pooled and pelleted and resuspended for co culture assays. For 48hr co culture, 1 x 10^4^ TAMs were plated with 5 x 10^4^ T cells in 200 μl of T cell complete media. After the incubation time, cells harvested were subjected to flow cytometric analysis assessing T cell activation and conditioned media harvested was used for IFN-γ ELISA as mentioned above.

### CCLE cell line expression analysis

mRNA expression data for *CREB* and *LIF* were obtained from the Cancer Cell Line Encyclopedia (CCLE) database via the Broad Institute’s portal (DepMap, https://depmap.org). Data corresponding to pancreatic ductal adenocarcinoma (PDAC) cell lines were extracted. The normalized expression values described in transcripts per million (TPM) for *CREB* and *LIF* were then imported into GraphPad Prism for correlation analysis. A linear regression analysis was performed, and the Pearson correlation coefficient (R) along with the p-value were reported to assess the strength and significance of the association in human PDAC cell lines.

### Lentiviral Transduction and Selection of CREB Overexpressing KPC Cells

Murine PDAC KPC cell line was engineered to overexpress CREB via lentiviral transduction Viral titer was generated in HEK293T cells using mammalian gene expression lentiviral vector (pLV-EGFP-EF1A-HA-HA-mCREB1; Vector ID-VB231120-1355wrv) collected after 48 and 72 hrs. pooled, clarified through a 40 μM filter and concentrated using Lenti-X Concentrator (Takara Bio, Cat.No # 631231) and resuspended in complete DMEM. KPC cells were seeded at a density of 2 × 10⁵ cells/well in 6-well plates and transduced with viral suspension. Following overnight incubation, the medium was replaced with fresh growth medium supplemented with 10% FBS. Transduced cells expressing EGFP were enriched by flow cytometric sorting and subsequently expanded for downstream applications.

## Acknowledgements

Grant Support: This study was supported by the R01 CA262526 grant from the National Cancer Institute (NCI) of the National Institutes of Health (NIH), the James Esther and King Biomedical Research Program (22K06) and Florida Cancer Innovation Fund (25C22) of the Florida Department of Health awarded to N.S. Nagathihalli. The Histopathology Core Service was conducted with the assistance of the Sylvester Comprehensive Cancer Center support grant, under the supervision of N. Nagathihalli. The research reported in this publication was supported by Sylvester Comprehensive Cancer Center and in part by the NCI of the NIH under Award Number P30 CA240139. The authors bear full responsibility for the content and the opinions expressed in this work, which may not necessarily reflect the official perspectives of the NIH.

The authors thank Dr. Michael VanSaun for his assistance with the genetic mouse models and Dr. Samara Singh for her help in the manuscript editing process. The authors would like to acknowledge the BioRender citations (see Supplementary Table). Research reported in this publication was performed in part at the Analytical Imaging Shared Resource (AISR), Onco-Genomics Shared Resource (OGSR;RRID:SCR_022502), Cancer Modeling Shared Resource (CMSR;RRID:SCR_022891), and Biostatistics and Bioinformatics Shared Resource (BBSR;RRID:SCR_022890) of the Sylvester Comprehensive Cancer Center at the University of Miami Miller School of Medicine, and in part by the National Cancer Institute of the National Institute of Health under Award Number P30-CA240139. We sincerely acknowledge the data generated for human PDAC by the Human Tumor Atlas Network (HTAN) WUSTL Atlas at the Siteman Cancer Center and the McDonell Genome Institute.

## Data Availability

All multi-omics datasets generated in this study have been deposited in the NCBI Sequence Read Archive (SRA) under the BioProject accession numbers PRJNA1363924, PRJNA1364047, PRJNA1364164, and PRJNA1364759.

