## Supplementary Figure Legends for "Targeting CREB remodels the immune microenvironment to enhance immunotherapy responses in pancreatic cancer"

**Supplementary Figure S1**. CREB expression in human and in the KPC GEMM of PDAC. **(A)** UMAP visualization of *CREB* mRNA expression across distinct cellular subtypes of HTAN pancreatic cancer dataset. **(B)** RNA-FISH combined with immunofluorescence (IF) analysis depicting *Creb* mRNA expression with CK-19 ducts within pancreatic tumor section from KPC GEMM. **(C)** Bar plot showing pancreas specific loss of *Creb* mRNA gene in *Creb* deleted *KPCC* $^{-/-}$ as compared to wild type *KPC* with intact Creb expression (*Left*). Bar plot showing preserved expression of *CREB* mRNA within the tail of *Creb* deleted *KPCC* $^{-/-}$ mice identical to wild type *KPC* mice (*Right*). **(D)** RNA-FISH combined with IF analysis depicting loss of *Creb* mRNA expression with CK-19 ducts within pancreatic tumor section from *KPCC* $^{-/-}$ vs. *KPC*. **(E)** Representative photomicrographs of segmentation strategy along with heat map of differentially expressed proteins in the pancreatic tumor sections from the *KPC* and *KPCC*$^{-/-}$ GEMM using NanoString DSP platform. **(F)** Representative photomicrographs depicting IHC based staining along with quantification of Ki67$^{+}$ cells within the pancreatic tumor sections of *KPC* and *KPCC*$^{-/-}$ mice (n=3 mice per group). *p<0.05; $^{ns}$ non-significant (p>0.05).

**Supplementary Figure S2**. Cancer cell intrinsic deletion of CREB in KPC PDAC. **(A)** Experimental schematic depicting the working strategy for generation and expansion of single clonal populations of CREB knockout (CREB$^{KO}$) tumor cells using CRISPR-Cas9 based genomic editing technology in KPC mouse derived PDAC tumor cell line (*Top*). Western blot depicting loss of total CREB protein expression in CREB$^{KO}$ as compared to KPC *Creb* wild type (CREB$^{WT}$) tumor cells (*Bottom*). (**B**) Bubble plot outlining the expression of canonical lineage cell cluster annotations in the scRNA seq conducted on CREB$^{WT}$ and CREB$^{KO}$ orthotopic mouse PDAC tumors using 10X genomics platform. **(C-D**) Violin plots showcasing relative fold change of mRNA transcripts expression (*CD206, Arg1, iNOS*) in bulk tumor lysates of CREB$^{KO}$ KPC orthotopic pancreatic tumors as compared to CREB$^{WT}$. (**E**) Violin plots showcasing relative fold change of mRNA transcripts expression (*Cxcl9, Cxcl10*) in bulk tumor lysates of the same biological experimental cohorts. (n=4-6 mice per group). (**F**) Detailed outline of the *ex-vivo* co culture experiment; TAMs isolated from CREB$^{WT}$ or CREB$^{KO}$ KPC orthotopic tumors (n=4 mice per cohort) were co-cultured for 48hr with splenic pan T cells from non-tumor bearing mice, followed by flow cytometric analysis of T cell activation and IFN- $\gamma$ ELISA secretion. Individual data points with mean ± SEM are shown and compared by two-tailed unpaired t test. *p<0.05; **p<0.01.

**Supplementary Figure S3.** CREB mediated regulation of LIF in PDAC. **(A)** Bar plots showing *Lif* gene expression via qPCR and **(B)** LIF secretion in the conditioned media via ELISA (pg/ml) from $CREB^{WT}$ and $CREB^{KO}$ KPC cells (n=4 technical replicates each). **(C)** Bar plots depicting increased LIF levels in conditioned media from CREB overexpressing KPC cells ($CREB^{OE}$) as compared to KPC cells expressing endogenous (wild type) CREB ($CREB^{NTV}$). **(D)** mRNA levels of *LIF* in normal pancreatic tissue (Normal; n=171) and primary PDAC patients (Tumor; n=179) from GTEx and PAAD TCGA database, respectively. **(E)** Spearman rank correlation analysis of mRNA levels between *LIF* and *CREB* in PDAC patients (PAAD TCGA) database. **(F)** Kaplan Meier survival curve analysis of combined gene signatures for *LIF* and *CREB* in PAAD TCGA database. **(G)** Spearman rank correlation analysis of mRNA levels between *LIF* and *CREB* in human PDAC cell lines using CCLE dataset extracted on DepMap portal. **(H)** qPCR based transcriptional profiling of *LIF* and *CREB* mRNA gene expression in human HPNE and PDAC cell lines. Individual data points with mean ± SEM are shown and compared by two-tailed unpaired t test. *p<0.05; ***p<0.001; ****p<0.0001.

**Supplementary Figure S4.** CREB regulated LIF mediates paracrine signaling to macrophages. **(A)** Schematic depiction differentiation of bone marrow derived myeloid progenitors into macrophages, followed by 48hr incubation with conditioned media from $CREB^{WT}$ or $CREB^{KO}$ cells and subjected to RNA sequencing (n=3 technical replicates each). **(B)** Heat map-based depiction of differentially expressed genes in bone marrow derived macrophages (BMDMs) in $CREB^{WT}$ vs. $CREB^{KO}$ conditions. **(C)** Schematic depiction of isolation of pan T cells from non-tumor bearing C57BL6 mice co cultured with BMDMs subjected to different experimental conditions to assess IFN- γ secretion via ELISA. **(D-E)** Quantitative representation of stacked bar plots illustrating IFN-γ secretion, measured by ELISA under multiple experimental conditions depicted in the figure. **(F)** Schematic representation of KPC tumor cells into the syngeneic C57BL/6 mice, followed by subcutaneous injection of LIFR blockade EC359 (15mg/kg) for 2 weeks of treatment. **(G)** Violin plots depicting mean fluorescence intensity (MFI) of Arg-1 and PD-L1 expression in Veh vs. EC359 treated pancreatic tumors (n=5-7 mice in each treatment arm). **(H)** ELISA-based validation of LIF depletion in the conditioned media from LIF knockout ($LIF^{KO}$)

KPC cell clones, generated using CRISPR-*Cas9* genome editing, compared with control (non-targeting vector [NTV] KPC cells (n=4 technical replicates each). **(I)** qPCR-based analysis of mRNA expression depicting the relative fold change of direct downstream targets of LIF signaling in $LIF^{NTV}$ vs. $LIF^{KO}$ KPC cell clones (n=3-4 technical replicates each). Individual data points with mean ± SEM are shown and compared by two-tailed unpaired t test. *p<0.05; **p<0.01,****p<0.0001; $^{ns}$ non-significant (p>0.05).

**Supplementary Figure S5.** Therapeutic targeting of CREB using the small-molecule inhibitor (CREBi) 666-15 in a KPC orthotopic model of PDAC. **(A)** Relative mRNA expression levels of *Lif* in vehicle (veh) and CREBi-treated bulk tumor lysates, n=4 mice in each group. **(B)** Violin plot depicting quantification of Mean Fluorescence Intensity (MFI) of Arg1, CD86 and PD-L1 *(left)* along with representative histogram plots in veh and CREBi treatment (*right*) (n=5-8, mice in each group). Individual data points with mean ± SEM are shown and compared by two-tailed unpaired t test. *p<0.05; **p<0.01,****p<0.0001.

**Supplementary Figure S6.** Gating strategy applied following flow cytometric analysis. **(A)** and **(B)** Flow cytometry gating and sub-gating strategy, beginning with live singlets, used to identify myeloid macrophages and T-cell subpopulations.

**Supplementary Figure S7.** Raw uncropped images of Western blot membranes for Supplementary Figure S2*A* **(A)** and Figure 4*D* **(B)**.
