## Supplementary Figures for "Targeting CREB remodels the immune microenvironment to enhance immunotherapy responses in pancreatic cancer"

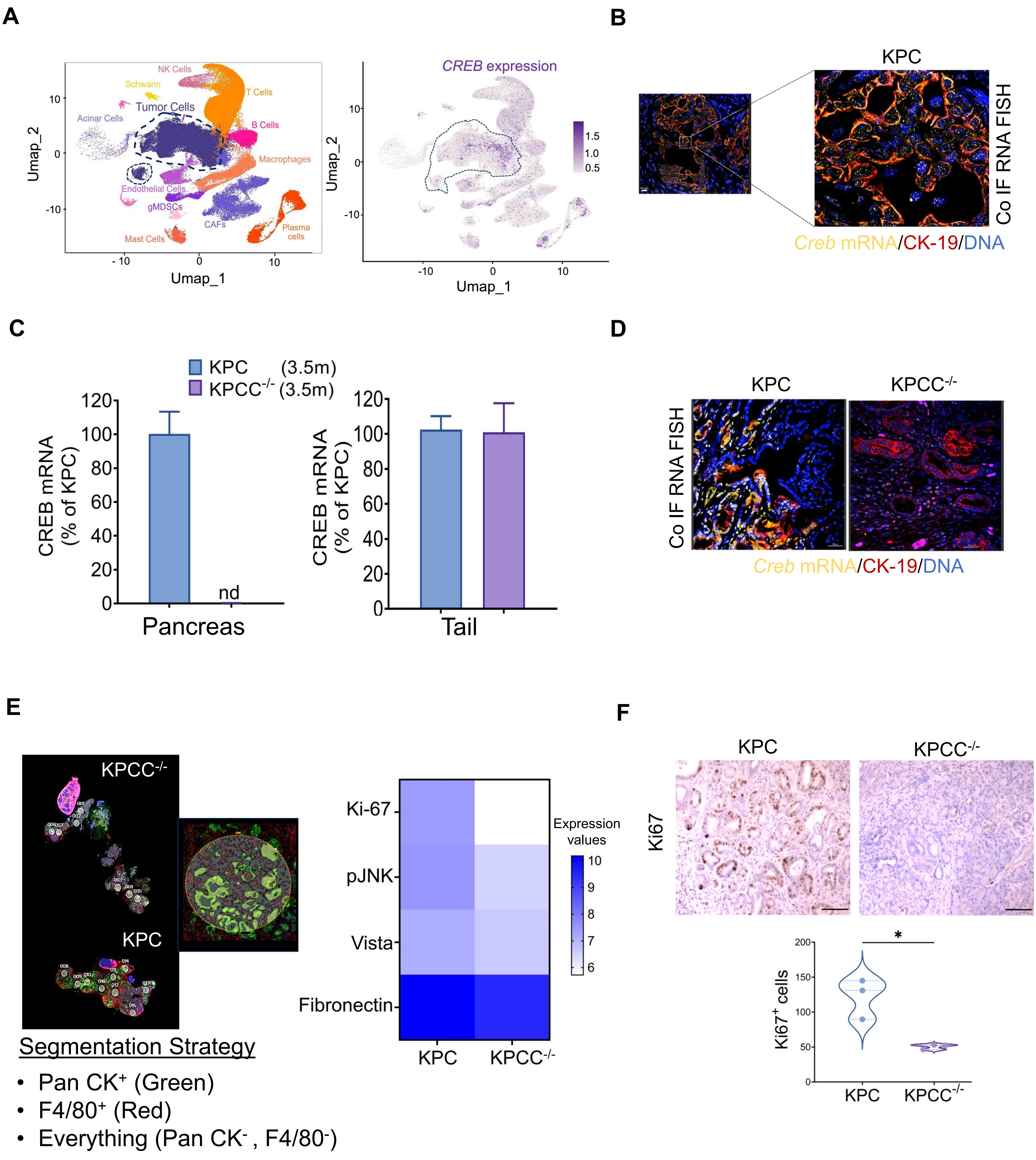


Supplementary Figure S1

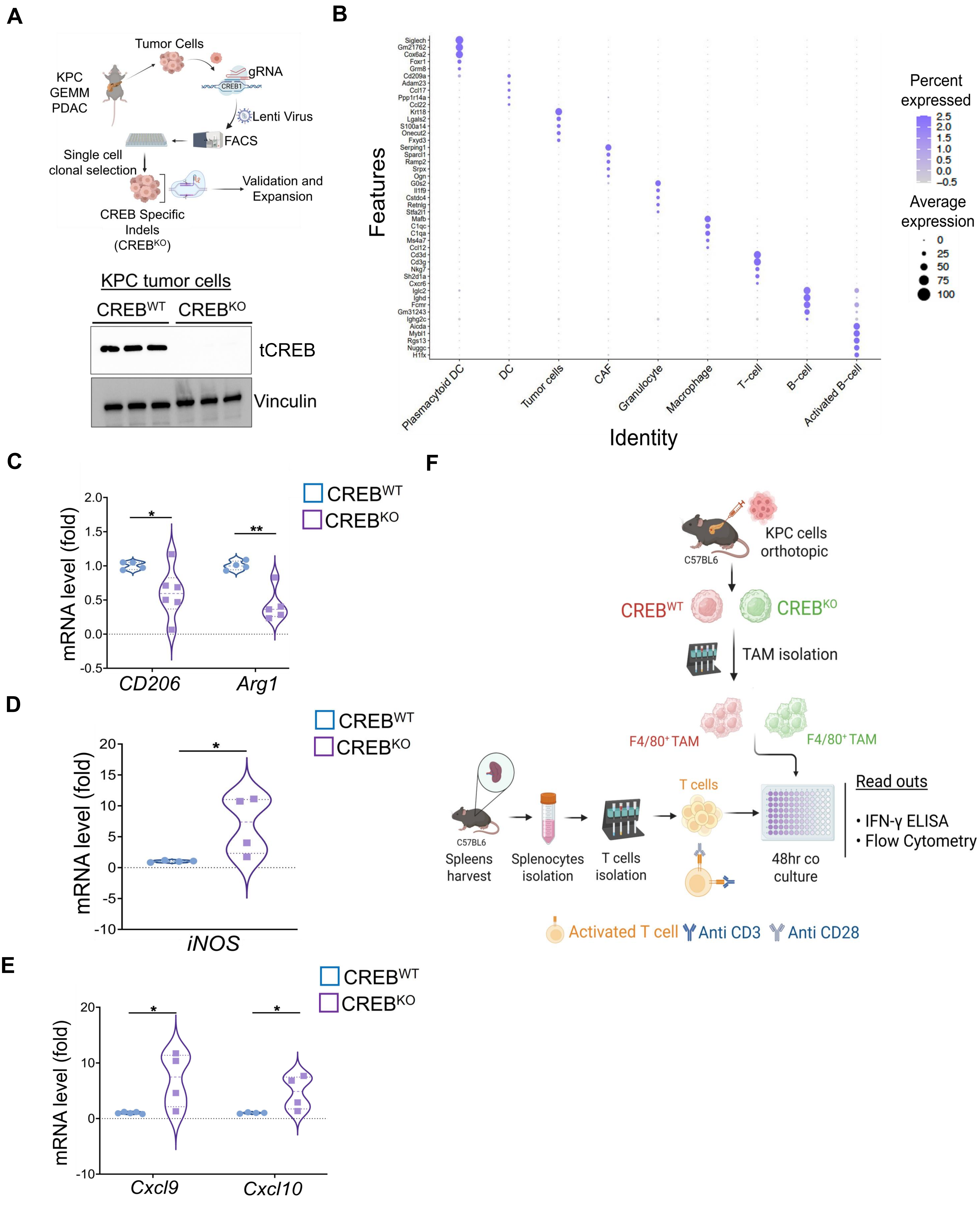


Supplementary Figure S2

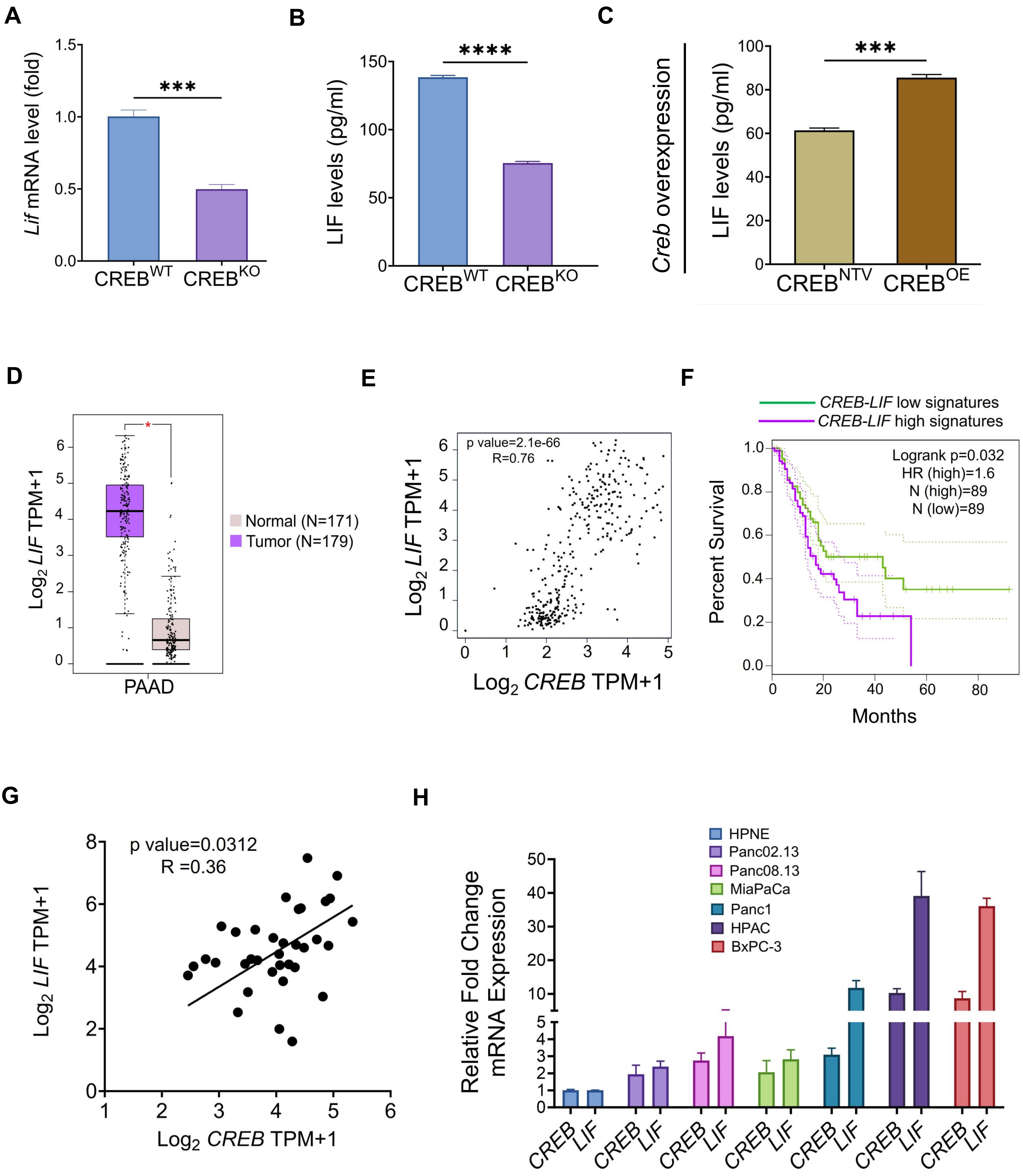


Supplementary Figure S3

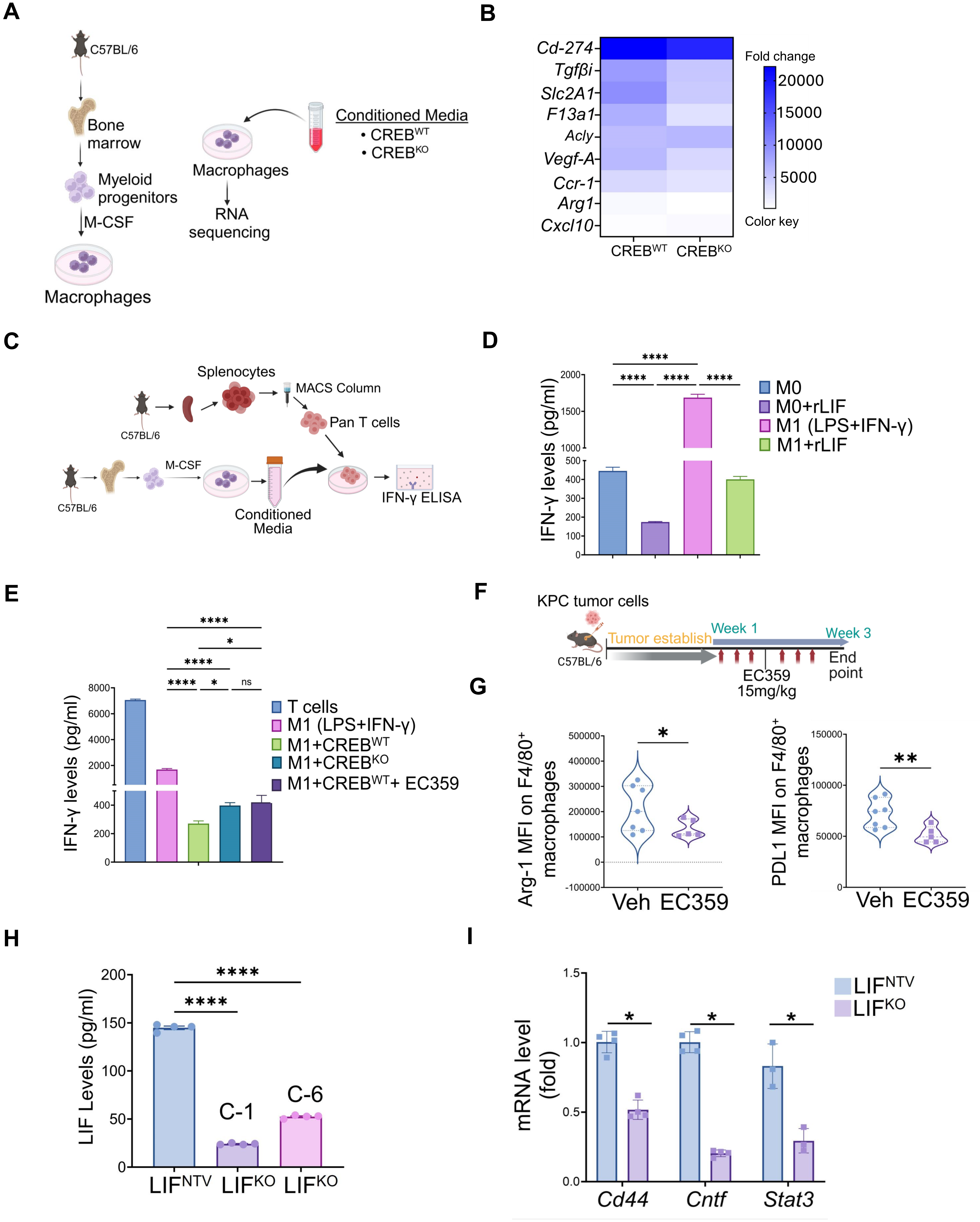


Supplementary Figure S4

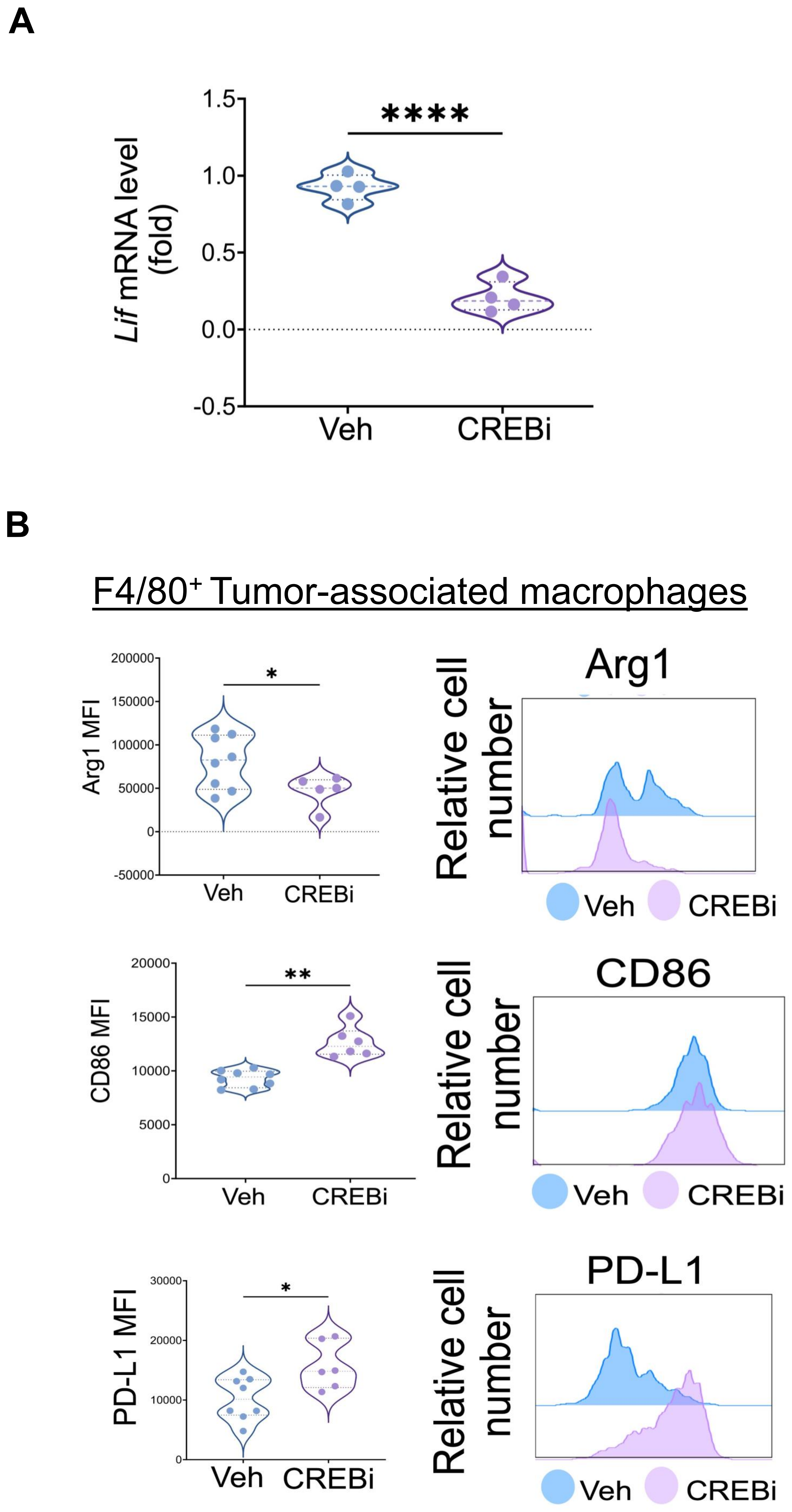


Supplementary Figure S5

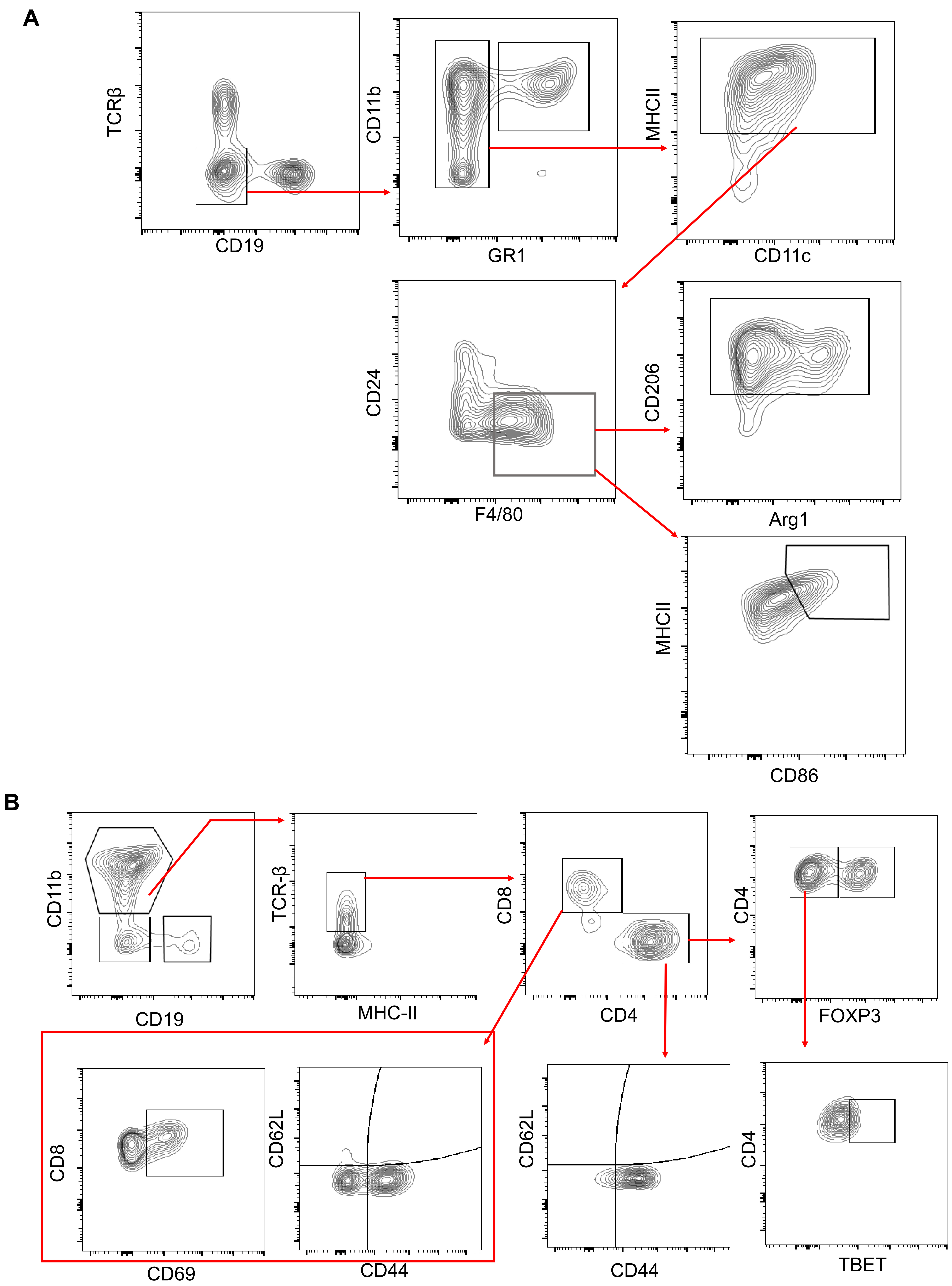


Supplementary Figure S6

**A**

Supplementary Figure S2A

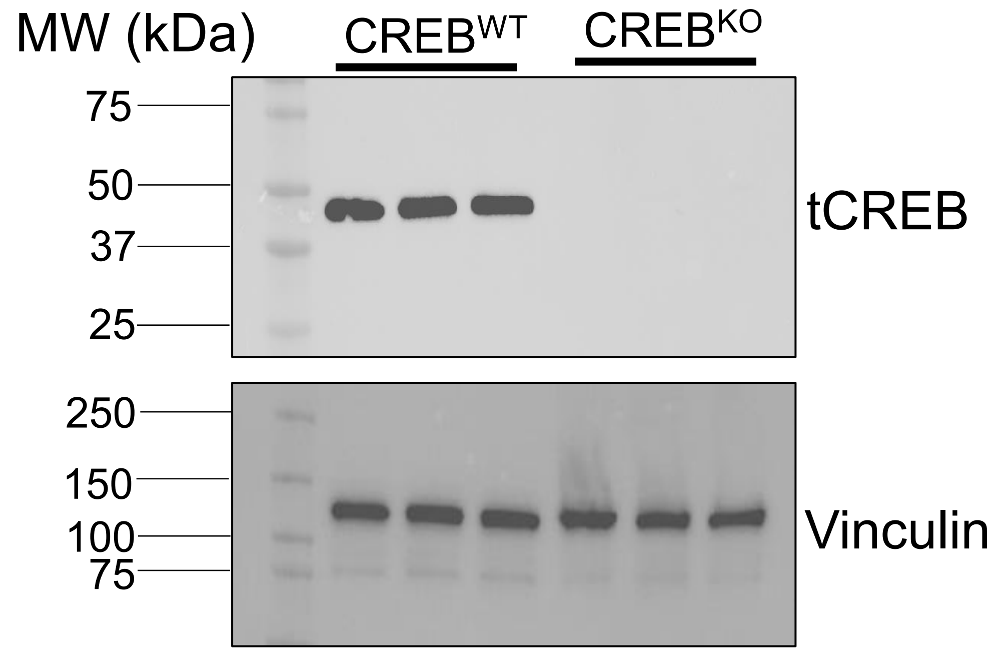


**B**

Figure 4D

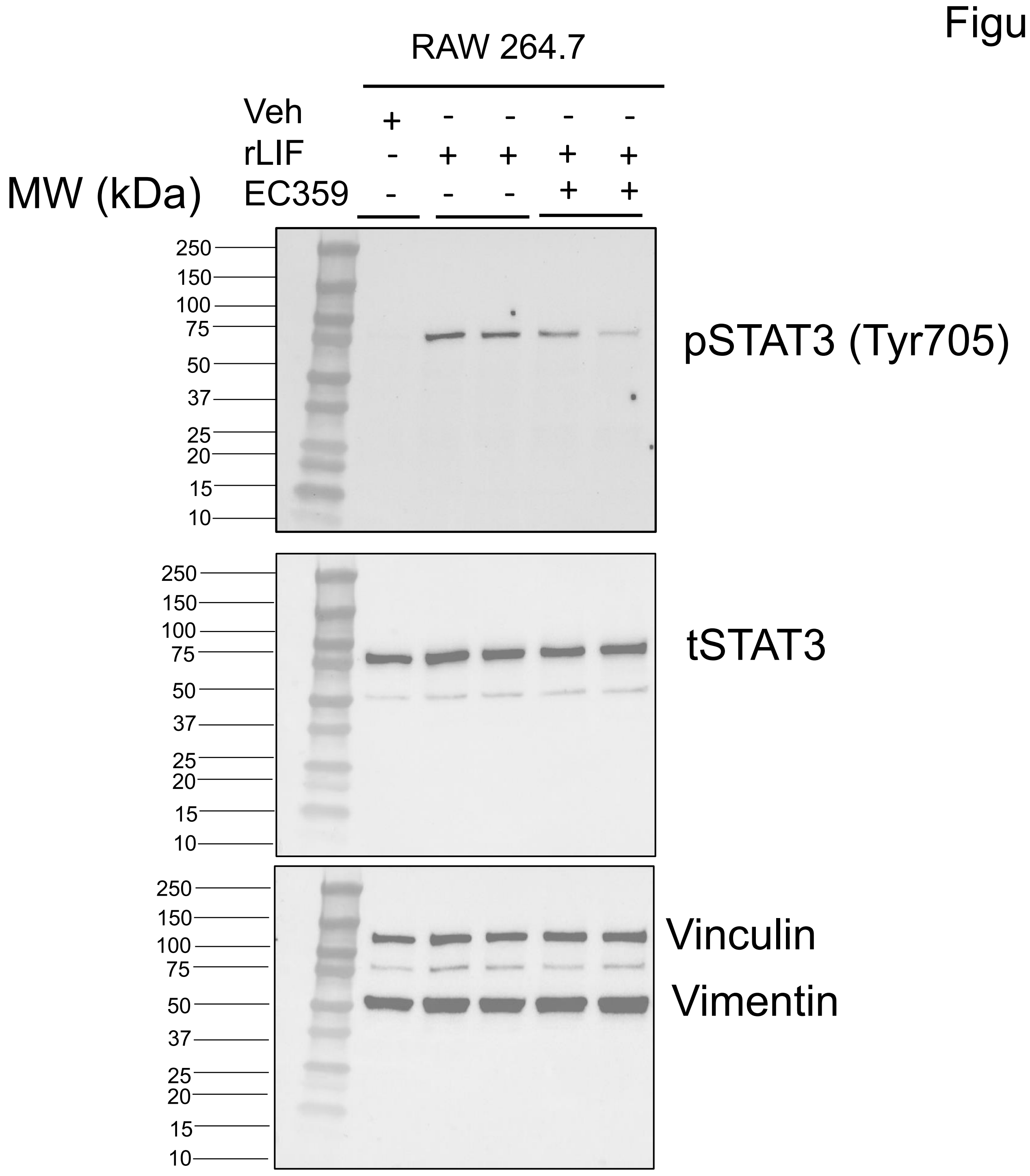


### Supplementary Figure S7
