## Supplementary Tables for "Targeting CREB remodels the immune microenvironment to enhance immunotherapy responses in pancreatic cancer"

| Figures | BioRender License Agreement Number |
|---|---|
| Fig. 1E | TT292D48PQ |
| Fig. 1I | TL292D54WI |
| Fig. 3A | XO292D5L2L |
| Fig. 3K | SW292D74XE |
| Fig. 4A | OQ292D7C50 |
| Fig. 5A | JG292D8U38 |
| Fig. 5B | GC292DAU76 |
| Fig. 5G | VL292DB5AS |
| Fig. 5K | IQ292DBDUD |
| Fig. 6A | VH292DBOAL |
| Fig. 6C | YZ292DBUHD |
| Supplementary Fig. S2A | EW292DCIFL |
| Supplementary Fig. S2F | AG292DIK1Q |
| Supplementary Fig. S4A | RJ292DIUNK |
| Supplementary Fig. S4C | ZM292DJ5LW |
| Supplementary Fig. S4F | DO292DJIC0 |

**Supplementary Table S1.** Primary antibodies for histological and Western blot analysis.

| Primary Antibodies | Supplier | Species | Catalogue number |
|---|---|---|---|
| pCREB (Ser-133) | Cell Signaling | Rabbit | 9198S |
| tCREB | Cell Signaling | Rabbit | 9179S |
| CK-19 | Abcam | Rabbit | Ab52625 |
| Ki67 | Cell Signaling | Rabbit | 12202S |
| CD68 | Cell Signaling | Rabbit | 76437T |
| F4/80 | Cell Signaling | Rabbit | 70076S |
| Arginase 1 | Cell Signaling | Rabbit | 93668S |
| CD206 | Cell Signaling | Rabbit | 24595S |
| Vinculin | Cell Signaling | Rabbit | 13901S |
| Vimentin | Cell Signaling | Rabbit | 5741S |
| LIF | Invitrogen | Rabbit | PA5-115510 |
| pSTAT3 (Tyr-705) | Cell Signaling | Rabbit | 9145S |
| tSTAT3 | Cell Signaling | Rabbit | 30835S |

**Supplementary Table S2.** Sequence of probes used for mice genotyping analysis.

| **Gene** | **Forward primer** | **Reverse primer** |
|---|---|---|
| Cre | GAA GGC ATT TGT GTA GGG TCA | GGC TGA GTG AGG GTT GTG AG |
| $Creb^{fl}$ | CTCTTCTTGCATCAAGCTTGGT | AGATCCCTCTAGGCATTTCTTCCT |
| $Creb^{WT}$ | CCAGTTACCTTCTAGGGAGCAGCTTACA | CAGGCCTGAGGTCTGGCTTCA |
| $Kras^{G12D}$ | GGCCTGCTGAAAATGACTGAGTATA | CTGTATCGTCAAGGCGCTCTT |
| $Tp53^{R172H/+}$ | CATCTACAAGAAGTCACAGCACATG | GGAGCAGCGCTCATGGT |

**Supplementary Table S3.** Primary antibodies for flow cytometry analysis.

| **Primary Antibodies** | **Fluorophore** | **Supplier** | **Species** | **Catalogue number** |
|---|---|---|---|---|
| CD45 | BUV-805 | BD Biosciences | Mouse | 741957 |
| TCR-β | BUV-661 | BD Biosciences | Mouse | 749914 |
| CD4 | PE-Cy7 | Biolegend | Mouse | 116016 |
| CD3 | PerCPCy5.5 | Biolegend | Mouse | 100218 |
| CD8 | BV-510 | Biolegend | Mouse | 100752 |
| CD69 | BUV395 | BD Biosciences | Mouse | 740220 |
| PD-1 | BV-605 | Biolegend | Mouse | 135220 |
| PD-L1 | BV-421 | Biolegend | Mouse | 124315 |
| CD44 | BUV-737 | BD Biosciences | Mouse | 612799 |
| CD62L | BUV-563 | BD | Mouse | 741230 |
| TBET | Kiravia Blue | Biolegend | Mouse | 644838 |
| FOXP3 | PE | eBiosciences | Mouse | 12-5773-82 |
| CD11b | BV-750 | Biolegend | Mouse | 101267 |
| F4/80 | AF-488 | Biolegend | Mouse | 123120 |
| Arg1 | APC | eBiosciences | Mouse | 17-3697-82 |
| CD206 | BV-650 | Biolegend | Mouse | 141723 |
| MHC-II | APC Fire-750 | Biolegend | Mouse | 107652 |
| CD-86 | BUV-737 | BD Biosciences | Mouse | 741737 |
| iNOS | PE-Cy7 | eBiosciences | Mouse | 12-5773-82 |
| Ly6C/Ly6G | BUV-563 | BD Biosciences | Mouse | 741226 |
| CD103 | BV-480 | BD Biosciences | Mouse | 566118 |
| Ly6G | PE-Cy5 | eBiosciences | Mouse | 15-9668-82 |
| CD19 | PE-Cy5.5 | eBiosciences | Mouse | 35-0193-82 |
| CD11c | AF-700 | Biolegend | Mouse | 117320 |
| CD24 | BV-605 | Biolegend | Mouse | 101827 |
| TIGIT | BV-421 | Biolegend | Mouse | 142111 |
| CD39 | AF-647 | Biolegend | Mouse | 143808 |
| LIF receptor (LIFR) | PE | Biolegend | Mouse | 158908 |
| Viability L/D Blue | UV excitation | Invitrogen | Mouse | L23105 |

**Supplementary Table S4.** Primers used in the study.

| **Mouse Primers** | **Company** | **Gene Globe-ID** |
|---|---|---|
| *Creb1* | Qiagen | PPM03382F-200 |
| *Lif* | Qiagen | PPM02988F-200 |
| *Mrc1* (CD206) | Qiagen | QT00103012 |
| *Arginase 1* (Arg1) | Qiagen | PPM31770C-200 |
| *Cxcl9* | Qiagen | PPM02973B-200 |
| *Cxcl10* | Qiagen | PPM02978E-200 |
| *iNOS (Nos2)* | Qiagen | PPM02928B-200 |
| *Cd44* | Qiagen | PPM03628F-200 |
| *Cntf* | Qiagen | PPM68695B-200 |
| *Stat3* | Qiagen | QT00148750 |
| *Rn18s* (18s ribosomal RNA) | Qiagen | PPM72041A-200 |
| **Human Primers** | **Company** | **Gene Globe-ID** |
| *CREB1* | Qiagen | PPH00808F-200 |
| *LIF* | Qiagen | PPH00813F-200 |
| *RN18S (18s ribosomal RNA)* | Qiagen | PPH84351A-200 |

**Supplementary Table S5.** Sequence of oligonucleotides used for mouse *Lif* LentiCRISPR genome editing

| Gene | Forward | Reverse |
|---|---|---|
| *Lif* gRNA | caccgCGGGACAGAGAAGACCAAGT | aaacACTTGGTCTTCTCTGTCCCGc |
| Non targeted control gRNA sequence | caccGGAAACGTTACATTCGACGC | aaacGCGTCGAATGTAACGTTTCC |
